# Cancer-keeping genes as therapeutic targets

**DOI:** 10.1101/2022.06.13.495906

**Authors:** Xizhe Zhang, Chunyu Pan, Xinru Wei, Meng Yu, Shuangjie Liu, Jun An, Jieping Yang, Baojun Wei, Wenjun Hao, Yang Yao, Yuyan Zhu, Weixiong Zhang

**Author notes:** Correspondence. XZ; YZ; WZ.

## Abstract

Finding cancer-driver genes – the genes whose mutations may transform normal cells into cancerous ones – remains a central theme of cancer research. We took a different perspective; instead of considering normal cells, we focused on cancerous cells and genes that maintained abnormal cell growth which we named *cancer-keeping genes* (CKGs). Intervention in CKGs may rectify aberrant cell growth so that they can be adopted as therapeutic targets for cancer treatment. We developed a novel approach to identifying CKGs by extending the well-established theory of network structural controllability, which aims at finding a control scheme (i.e., a minimal set of non-overlapping control paths covering all nodes) and control nodes (driver genes) that can steer the cell from any state to the designated state. Going beyond driver genes defined by one control scheme, we introduced *control-hub* genes located in the middle of a control path of *every* control scheme. Control hubs are essential for maintaining cancerous states and thus can be taken as CKGs. We applied our CKG-based approach to bladder cancer (BLCA) as a case study. All the genes on the cell cycle and p53 pathways in BLCA were identified as CKGs, showing the importance of these genes in cancer and demonstrating the power of our new method. Furthermore, sensitive CKGs that could be easily changed by structural perturbation were better suited as therapeutic targets. Six sensitive CKGs (RPS6KA3, FGFR3, N-cadherin (CDH2), EP300, caspase-1, and FN1) were subjected to small-interferencing-RNA knockdown in two BLCA cell lines to validate their cancer-suppressing effects. Knocking down RPS6KA3 in a mouse model of BLCA significantly inhibited the growth of tumor xenografts in mice. Combined, our results demonstrated the value of CKGs as therapeutic targets for cancer therapy.

**Key points:**

- Focus on genes that maintain aberrant cell growth, named *cancer-keeping genes* (CKGs).
- Develop a novel approach for finding CKGs by extending the well-estabilished theory of network structural controllability to total network controllability.
- Apply the new method to bladder cancer and experimentally validated the cancer-suppressing function of six CKGs in two bladder cancer cell lines and that of one CKG in bladder cancer mice.

## INTRODUCTION

A primary objective of cancer research is to identify genes that may trigger tumorigenesis or promote aberrant cell growth, which are referred to as cancer-driver genes (CDGs)^1, 2^. CDGs can help elucidate cancer etiology^3^ and be used as diagnostic biomarkers and therapeutic targets^4^. Most existing methods for CDG discovery look for genes with substantial mutations that can separate cancer subjects and normal controls^5, 6, 7^. Despite many efforts^1, 2, 8^, CDGs that can be discovered seem to be saturated with increasing sample sizes^9^.

Deviated from the prevalent mutation-based methods for CDG finding is a recent approach based on network structural controllability^10, 11^. Structural controllability specifies how a networked system can be driven from any state to the desired state in finite time by exerting stimuli on the driver nodes defined by a control scheme (i.e., a minimal set of non-overlapping paths covering all nodes) for the network^10^. Network controllability has been applied to various biological networks^12–14^, including protein-protein interaction networks^15^, gene regulatory networks^16, 17^, and metabolic networks^18, 19^. Instead of analyzing individual genes in isolation, this approach organizes all genes of interest in a network and analyzes them all together to identify driver nodes or genes. When applied to cancer regulatory networks, it identifies driver nodes that are regarded as CDGs and therapeutic targets for cancer treatment^20^.

However, two critical issues hinder network controllability from becoming a practical method for finding CDGs. Firstly, given a network, the control scheme is often not unique, but rather numerous control schemes and different sets of driver nodes exist^21, 22^. It is difficult, if not infeasible, to determine the most effective control scheme for the network. All control schemes may be compared to select the best, e.g., one with the shortest control trajectory to the desired state. However, finding all control schemes, a #P-hard problem^23^, is computationally prohibitive. In addition, one control scheme may contain a substantial number of driver genes^21^ which may need to be stimulated together to control the cell. It is difficult to manipulate many genes at once for disease treatment. Secondly, a fundamental but mostly neglected assumption underlying structural controllability methods is that the biological network being analyzed is a model describing both cancerous and normal states of the cell so that mutations in some genes can drive the cell from a normal state to a cancerous state, resulting in cancer. However, little is known about which states are normal and which other states are cancerous, so a driver node in the network may not necessarily be a CDG.

We adopt a different perspective on cancer and cancer treatment. Instead of considering mutated genes that may transform normal cells into abnormal ones, we are interested in genes that maintain cancerous cell states, which we call *cancer-keeping genes* (CKGs). We assume that intervention to such genes may terminate or prevent aberrant cell differentiation and proliferation and hypothesize that CKGs are ideal candidate biomarkers for diagnosis and therapeutic targets for therapy. To this end, we focused on homogenous networks that model cancer cells and utilized and extended the theory of structural controllability. Importantly, we went beyond one control scheme and considered the overall structural controllability governed by all control schemes of a network. We introduced *control hubs* located in the middle of a control path of *every* control scheme. We developed a polynomial-time algorithm for finding all control hubs without computing all control schemes. We were particularly interested in those control hubs that were sensitive to changes to network structures so better suited as therapeutic targets. In a case study, we applied our approach to bladder cancer (BLCA) and identified 35 sensitive CKGs for the disease. One important finding was that the genes on the cell cycle and p53 pathways were sensitive CKGs. This result showed that these genes were indeed critical for maintaining BLCA and demonstrated the power of our new method. Using small-interferencing RNA (siRNA) knockdown, we experimentally validated six sensitive CKGs *in vitro* by examining their effects on the proliferation of two BLCA cell lines. We also validated RPS6KA3, the only sensitive CKG on the p53 pathway, *in vivo* in a mouse model of BLCA by showing its prohibitive function on tumor progression.

## RESULTS

The new CKG-based approach consists of four major steps: 1) Construction of a biological network describing gene-gene interactions/associations for the disease of interest; 2) Identify control hubs in the network and from which, identify CKGs; 3) Analyze the functions of selected CKGs; and 4) Experimental characterize the cancer suppression functions of sensitive CKGs in cell lines and or animal models. In the sequel, we present these major components along with the rationale and key ideas of the new approach. We focus on BLCA as an application, but the new method is general and applicable to other cancers.

### Construction of bladder cancer gene regulatory network

We developed a novel method for constructing a gene regulatory network for cancer study, specifically studying BLCA (Figure 1A). The network consists of cancer-related genes and interactions of the ten most important and common types of interactions and signaling pathways (Supplemental eTable 1) from five well-curated, disease- and cancer-related pathway databases, including the NCI Pathway Interaction Database^24^, PhosphoSite Kinase-substrate information^25^, HumanCyc^26^, the Reactome^27^, and PANTHER Pathways^28^ (see Materials). These databases were created in part by disease studies, so they contain information on mutated genes and disease- and cancer-related pathways. For example, Reactome consists of the KEGG databases that include a disease database and a drug database. Phosphosite contains data on missense mutations from UniProtKB, TCGA, and many other sources, which are collected from more than 2,000 diseases and syndromes, and has data on polymorphisms associated with hundreds of cancers.

**Figure 1.**
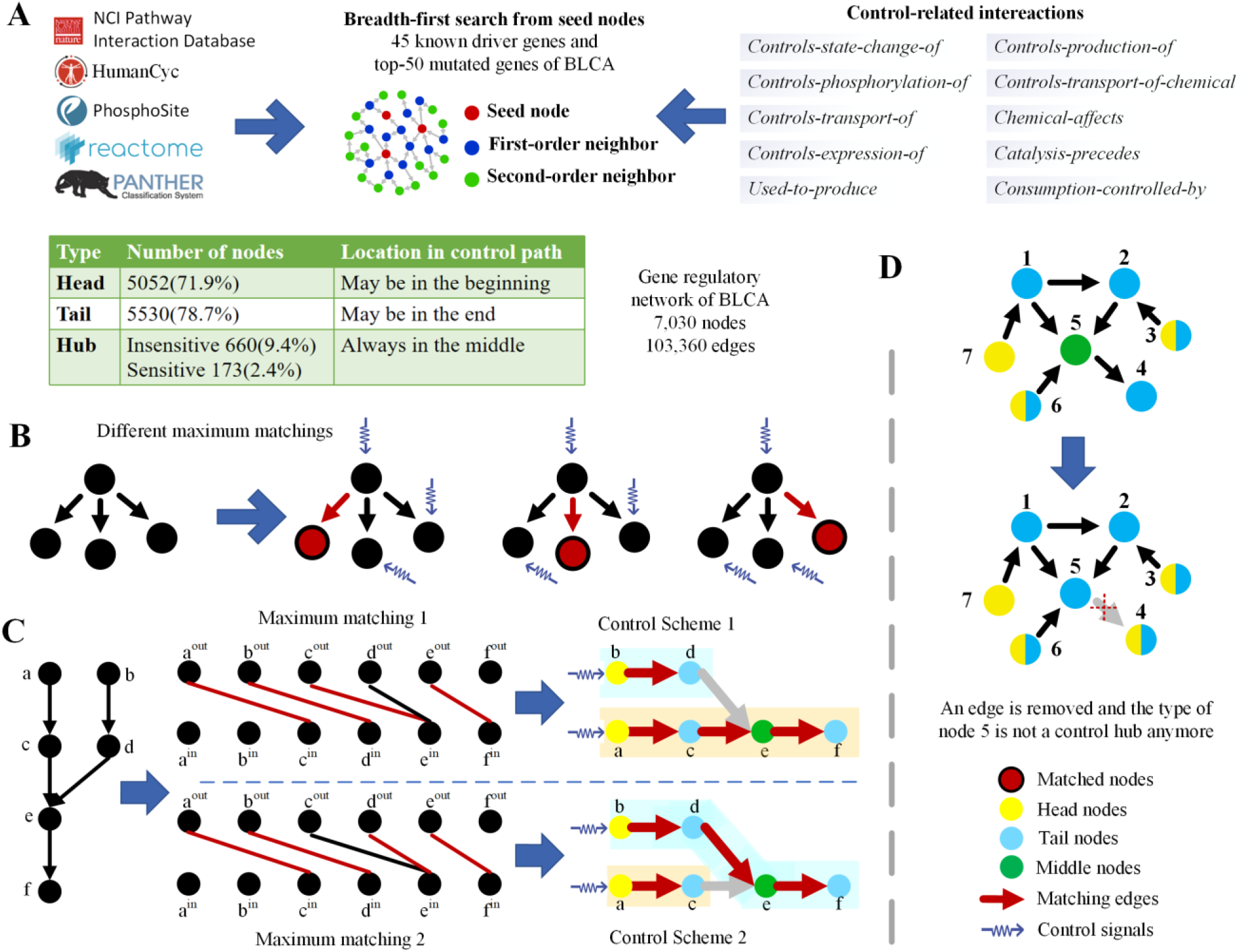
BLCA gene regulatory network construction and control hubs of complex networks. **A)** Construction of a BLCA gene regulatory network, BLCA_GRN. Ten types of control-related gene/protein interactions were extracted from five high-quality pathway databases. A breadth-first search was then performed, starting from 45 known BLCA driver genes and the 50 most highly mutated genes of BLCA. The traversed nodes and edges constitute BLCA_GRN. **B)** An example of a simple network with three different maximum matchings. The unmatched nodes are driver nodes, and the red edges are matched edges. **C)** Two different control schemes of a simple network. A node may lie at the head, tail, or middle of a control path. Node *e* resides in the middle of some control paths of the two control schemes, becoming a control hub of the network. **D)** Sensitive control hubs of a simple network. After removing edge *e*(5,4), node 5 changes from a control hub to a tail node, therefore, it is a sensitive control hub of the network.

To construct a BLCA regulatory network, we used some cancer driver genes as seeds to traverse the ten disease- and cancer-related interaction databases. From the data of TCGA Research Network (https://www.cancer.gov/tcga), we chose 45 known BLCA cancer-driver genes^29^ and 50 most mutated genes in BLCA as seed nodes (Supplemental eTable 2). A breadth-first search starting from the 95 seed genes was used to traverse the ten major types of control-related interactions in the databases. The traversed genes and the interactions formed the BLCA gene regulatory network (BLCA_GRN) with 7,030 nodes (genes) and 103,360 directed edges (Figure 1A; see Supplemental File 1).

### Control hub nodes of a network

Our new method hinges upon the concept of control hubs of a network. They are the most vulnerable spots for structural controllability, so intervention on any of them may render the network not controllable by stimuli to the network. When applied to network models of cancer cells, interventions on control hubs may terminate aberrant cell differentiation and proliferation, thus they may be taken as therapeutic targets.

A key observation supporting structural controllability is that a node in a directed network can control one of its outgoing neighbors^30^ and the controlled neighbor can control a neighbor of its own and so forth. These nodes collectively form a *control path* that has a head node at the beginning of the path, also referred to as *control node*^12^ or *driver node*^11^, a tail node at the end of the path, and middle nodes if any (Figures 1B and 1C, Supplemental eMethods 1 and 2). The set of the smallest number of control paths and their driver nodes constitute a *control scheme* of the network (Figure 1C). Importantly, the control scheme is not unique for a complex network^21, 22^. A node may take different positions and thus play distinct roles in different control schemes (e.g., nodes *c* and *d* in Figure 1C). It is difficult to determine the best control scheme. We may compute all control schemes to select the best. However, it is computationally prohibitive to derive all control schemes because the problem is #P-hard^23^, meaning that no polynomial algorithm is known for the problem. Furthermore, given all control schemes, it is nontrivial to determine the best because little is known about the network dynamics and different optimality criteria (e.g., the fewest nodes to be controlled versus the fewest steps to reach the desirable state) will lead to distinct control schemes. Moreover, a substantial number of nodes may serve as driver nodes in different control schemes^21^, making it costly to control the network. All these unfavorable factors cast doubts on the utility of structural controllability for finding cancer driver genes.

We go beyond one control scheme and consider the overall structural controllability of a network by considering all control schemes. To ameliorate the computational burden, we avoid computing all control schemes and focus on all *control hubs* that are middle nodes of some control paths of *every* control scheme (e.g., node *e* in Figure 1C). An eminent feature of the control hub is that it is essential for maintaining the overall structural controllability of the network – perturbation to or blockade of any control hub may void all control schemes and consequently take apart the overall controllability of the network. Therefore, it is critical to protect all control hubs to maintain structural controllability and the validity of the network control model. This implies that control hubs are the most vulnerable spots for structural controllability. Here, we explicitly explore and exploit this property of control hubs.

However, identifying all control hubs of a large network is technically nontrivial. We developed a novel polynomial-time algorithm for finding all control hubs without computing all control schemes^31^. The algorithm is based on previous work on enumerating all driver nodes^21, 32^. It first identifies the head and tail nodes of the control paths of all control schemes and subsequently identifies control hubs. It has a complexity of *O*(*n×m*) on a network with *n* nodes and *m* edges.

### Sensitive Control hub nodes as cancer-keeping genes

We consider a model representing cancerous cells and focus on the overall network controllability using control hubs. When a structural change turns a control hub into a non-control hub, the network is no longer controllable by any control scheme. As a result, the cell may transition out of the current cancerous state to, presumably, the normal state. Therefore, we name control hub genes in a cancer regulatory network *cancer-keeping genes* (CKGs) because they maintain the network controllability of the model.

We expect that some control hubs are more sensitive and vulnerable to external perturbations than others and may be better therapeutic targets. We are particularly interested in those control hubs that can be turned into non-control hubs when a single edge is removed from the network as a perturbation (Figure 1D), which we call *sensitive control hubs* or *sensitive CKGs* (sCKGs). All sCKGs can be identified by removing every edge of the network one at a time (see Methods).

### Cancer-keeping genes are essential in the gene regulatory network of BLCA

We applied our novel CKG approach to the BLCA gene regulatory network BLCA_GRN containing 7,030 genes and 103,360 interactions (Figures 1A and 2A, Supplemental File 1). One control scheme has 3,115 driver genes (44.3% of all 7,030 genes) and there exist 5,052 driver genes (71.9%) for all control schemes (Figure 2B). So many control schemes and nearly two-thirds of all genes being driver genes made it infeasible to choose the right control scheme for BLCA when applying structural controllability.

**Figure 2.**
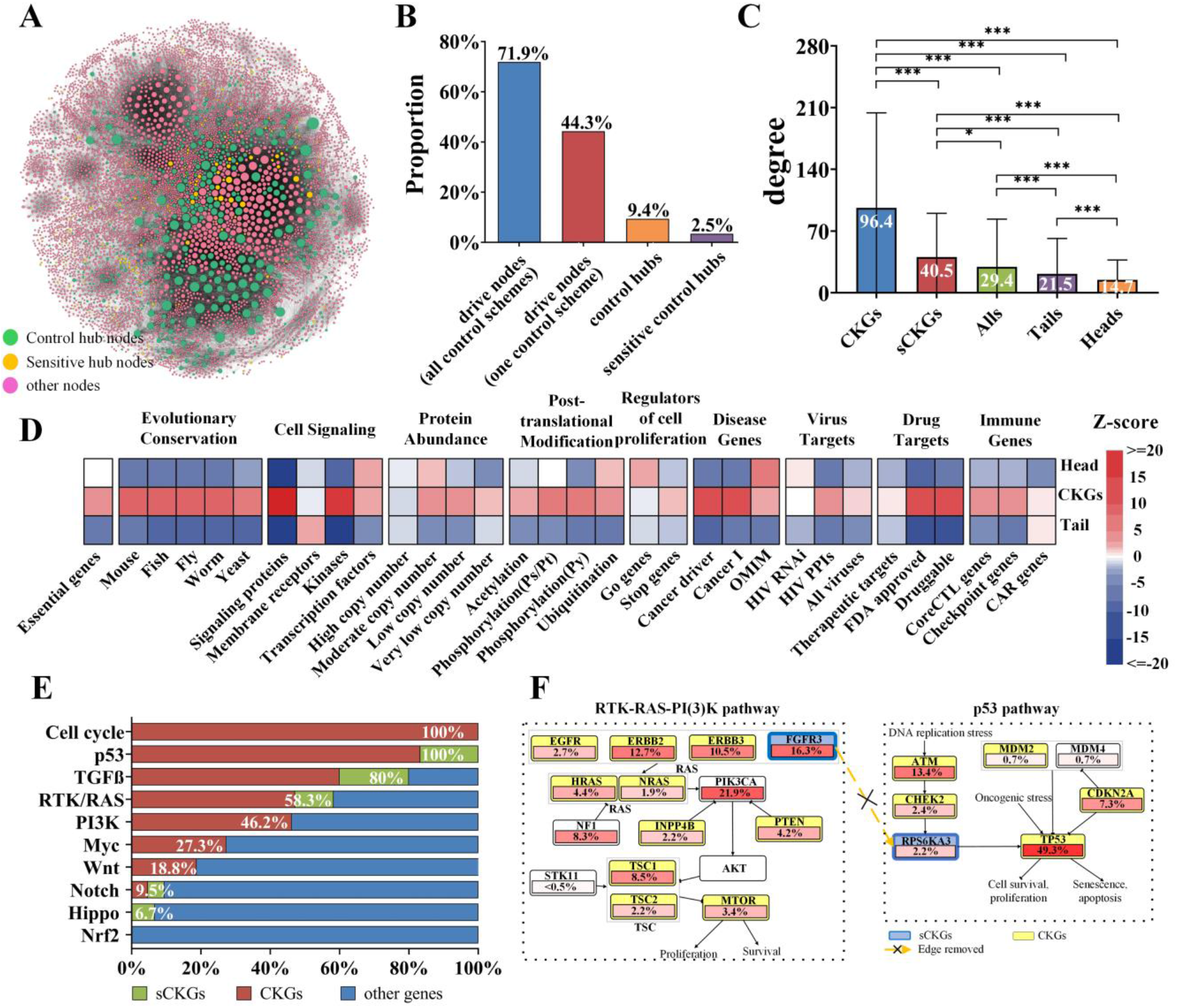
Control hubs or cancer-keeping genes (CKGs) in the gene regulatory network of bladder cancer (BLCA_GRN). **A)** Topological structure of BLCA_GRN. Node size is proportional to node degree. The network contains 7,030 genes and 103,360 directed interactions. **B)** Proportions of the driver nodes, control hubs, and sensitive control hubs in BLCA_GRN. **C)** Node degrees of different types of genes in BLCA_GRN. Control hubs have a significantly higher average degree than the other genes. One-way ANOVA (analysis of variance) was adopted, followed by multiple comparison tests (post-hoc) with a 0.05 significance level. ***p < 0.001; **p < 0.01; *p < 0.05. **D)** Control hubs of BLCA_GRN are enriched in the context of essentiality, evolutionary conservation, cell signaling, protein abundance, post-translational modifications (PTMs), regulators of cell proliferation, diseases, virus targets, drug targets, and immune regulation. **E**) Proportions of sensitive CKGs and CKGs in ten key cancer signaling pathways with significant genetic variations. Most of the genes in these pathways are CKGs. **F**) Two TCGA-analyzed signaling pathways of BLCA. FGFR3 and RPS6KA3, two control hubs, reside upstream of the most mutated genes in the pathways. The removal of edges from FGFR3 to RPS6KA3 will change their node types in the control scheme and therefore, make them sensitive CKGs.

Our CKG approach identified 660 CKGs in BLCA_GRN, which are 9.4% of all 7,030 genes and 13.1% of all 5,052 driver genes in the network (Figure 2B). The 660 CKGs have several characteristics. First, they have greater connectivity in the network, with an average degree of 96.4, than the other nodes which have an average degree of 29.4 (Figure 2C), reflecting the greater controlling power of the CKGs. Next, we characterized CKGs, head and tail genes in the context of essentiality, evolutionary conservation, and regulation at the levels of translational and posttranslational modifications (PTMs) as has been done previously^33^. The results showed that the CKGs were significantly enriched in most of the above datasets (Figure 2D, Supplemental eFigures 2-4). Interestingly, the CKGs were significantly enriched with pathogenic mutants and targets of viruses, drugs, and immunotherapies (Figure 2D, Supplemental eFigures 5 and 6). Altogether, these results revealed that CKGs play crucial regulatory roles.

Comparing the potential targets of tumor immunotherapy including core cancer-intrinsic CTL-evasion genes (coreCTL)^34^, CAR therapy targets^35^, and immune checkpoints^36, 37^ showed that 27 CKGs were known targets of tumor immunotherapy, in which 20 CKGs were coreCTL genes and 7 CKGs were CAR genes or resided at immune checkpoints. Surprisingly, although 437 CKGs were not directly immune-related, their direct neighbors in the network were immune regulatory genes (Supplemental File 2). Combined, these results suggested that the 660 CKGs were involved in multiple regulatory processes of tumor immunity, reflecting their functional importance.

To assess the potential oncogenic functions of the 660 CKGs, we examined their involvement in ten key cancer-related signaling pathways that have genes with extensive genetic variations in 33 types of cancer as analyzed by TCGA^38^. Interestingly, 70.4% of the genes in four cancer signaling pathways (i.e., the cell-cycle, p53, TGFß, and RTK-RAS pathways) were CKGs (Figures 2E and 2F, Supplemental eTable 4). Remarkably, all the genes in the cell-cycle and p53 pathways were CKGs (Figure 2E), which are key signaling pathways for cell differentiation and proliferation. Many CKGs (e.g., ACVR1B in the TGFß pathway and FGFR3, KIT, NTRK2, and BRAF in the RTK-RAS pathway) were located upstream of the cell-cycle and p53 pathways. Compared to nine popular node/gene identification methods (Supplemental eFigure 7), such as that based on node connectivity and centrality, more CKGs are in the cell-cycle, p53, TGF β, and Myc pathways, which are all known to regulate cell differentiation and proliferation^38^. These results indicated that CKGs were tightly regulated and played critical roles in cancer, particularly BLCA.

### Sensitive cancer-keeping genes have oncogenic functions and clinical importance in BLCA

To select a small subset of the 660 CKGs as potentially druggable targets^39^ for treating BLCA, we looked for sensitive CKGs (sCKGs) in BLCA_GRN. The rationale for edge removal as network perturbation is that many cancer drugs are kinase receptor inhibitors, thus blocking protein-protein interactions and acting as edge removal. One example is an irreversible fibroblast growth factor inhibitor Futibatinib^40^ for BLCA. We name an edge *sensitive edge* if removing it renders a CKG to become an sCKG. An sCKG has at least one sensitive edge associated; the more sensitive edges that an sCKG is associated with, the more sensitive and druggable it is. Of the 660 CKGs, 173 (26.2%) were sCKGs, among which more than 35.8% were associated with more than one sensitive edge (Figure 3A). All sensitive edges had high confidence scores^41^ (Supplemental eTable 5), indicating that highly confidently the sCKGs were sensitive to such external stimuli. The 173 sCKGs have several characteristics. They predominantly had moderate or low connectivity; their average connectivity was less than half of that of the other CKGs (Figure 2C). Counter-intuitively, the sCKGs had significantly lower rates of genetic alteration (including mutation, copy number variation, or homozygous deletion) than their neighbors in BLCA_GRN (Figure 3B, Supplemental File 2), so they may not be detected by conventional frequency-based^42, 43^ and network-based methods^42, 44^.

**Figure 3.**
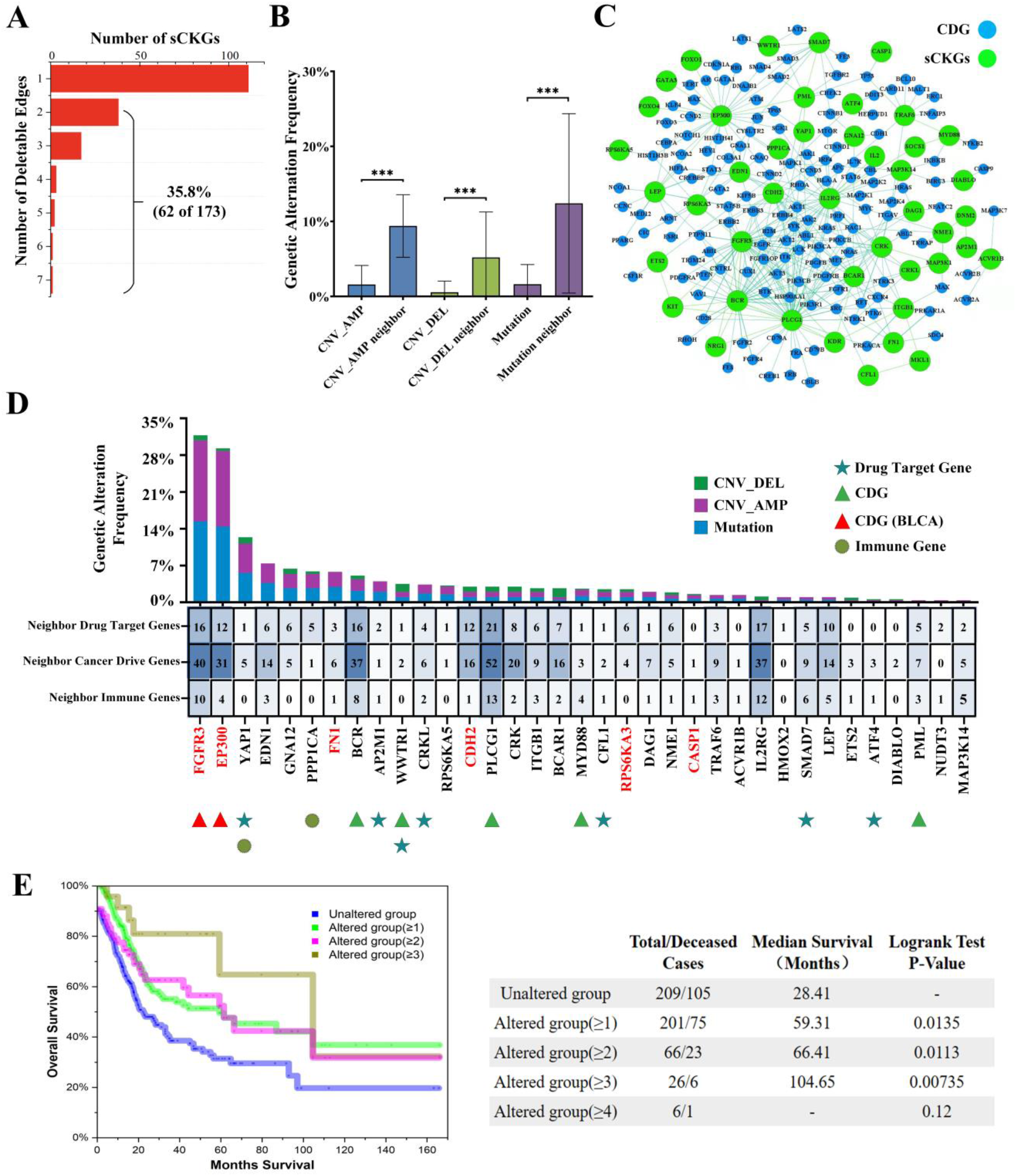
Characterizing sensitive CKGs (sCKGs) in bladder cancer gene regulatory network. **A)** The distribution of the number of edges that could change an sCKG. Most sCKGs have less than three edges that may change their node types, indicating they are robust to random structural perturbations. **B)** Differential analysis of the mutation frequencies between the sCKGs and their neighbors. The mutation frequencies of sCKGs are significantly lower than their neighbors for the sample from the cBioPortal database. **C)** The subnetwork of sCKGs and their surrounding cancer driver genes. **D)** Characterizing of 35 sensitive CKGs. Twenty-eight sCKGs are not cancer driver genes and the rest 7 are. Most of these CKGs are directly connected to drug targets, cancer driver genes, and/or immune genes in BLCA_GRN. The CKGs with names in red are experimentally studied (Figure 4). **E)** Survival analysis of the 35 sensitive CKGs in BLCA. Simultaneous alterations to more than one CKG will significantly increase the survival rate of patients from 59.31 months (one sCKG altered) to 104.65 months (three or more sCKGs altered).

We mapped the 173 sCKGs to the indispensable genes of the human PPI network^33^ resulting in 35 sCKGs that might be better therapeutic targets (Figure 3D). Most of these sCKGs were directly connected to the known cancer driver genes^29^, cancer therapeutic targets^45^ and/or immune genes^35–37^ in BLCA_GRN (Figures 3C and 3D, Supplemental eFigure 8) and were enriched with regulatory genes (Supplemental eFigure 9). For example, 26 of the 35 sCKG had cancer driver genes as direct neighbors (Figure 3D, Supplemental eTable 6), 30 (85.7%) were directly connected to cancer therapeutic targets (Figure 3D, Supplemental eFigure 10A, and eTable 7), and 27 (77.1%) were directly connected to immune genes (Figure 3D, Supplemental eFigure 10B, and eTable 8).

Remarkably, many of the 35 sCKGs had beneficial epistatic relations. Increased mutations to more than one of these genes significantly increased the overall survival rates of many BLCA subjects based on the TCGA PanCancer clinical data (see Methods). Comparing the median survival times of all BLCA patients with two or more of the 35 sCKGs being mutated revealed that the median survival time would increase from 28.41 months (for the unaltered group) to 59.31 months (for the one sCKG altered group), 66.41 months (for the two sCKGs altered group), and 104.65 months (for the three and more sCKGs altered group) under the stringent criterion of the Logrank Test p-value <0.015 (Figure 3E). This result strongly suggested that the sCKGs were excellent diagnostic and prognostic biomarkers and therapeutic targets for BLCA.

Two sCKGs (FGFR3 and RPS6KA3) deserved further scrutiny. FGFR3 is a well-characterized CDG for BLCA^46^ and FGFR3 and RPS6KA3 are involved in two key pathways of BLCA, RTK-RAS-PI(3)K and p53^47^ (Figure 2F). Remarkably, FGFR3 is upstream of PIK3CA in the RTK-RAS-PI(3)K pathway, and RPS6KA3 is upstream of TP53 in the p53 pathway, suggesting that FGFR3 and RPS6KA3 are upstream drivers of two key CDGs. Furthermore, RPS6KA3 is a substrate of FGFR3 so they are related and share many functions. FGFR3 can phosphorylate Y529 and Y707 of RPS6KA3 to assist ERK1/2 to connect to RPS6KA3 and keep RPS6KA3 active^48^. Note that the interaction between FGFR3 and RPS6KA3 was critical. The removal of the link between the two genes in BLCA_GRN would change the two genes from control hubs to head nodes and subsequently invalidate all control schemes for BLCA_GRN, suggesting that the two genes must play critical roles in BLCA.

Immunotherapy has been adopted in treating BLCA and several drugs (e.g., Atezolizumab and Avelumab) have been developed to target immune checkpoint inhibitors (e.g., PD-1 and PD-L1)^49^. Among the six well-known genes targeted by BLCA drugs, five (CD274/PD-L1, CTLA4, IL12B, PTGS2, and TFDP1) were head or tail nodes except FKBP1A (Supplemental eTable 9). Nevertheless, all six genes were well connected with CKGs (i.e., control hubs) in BLCA_GRN. For example, CTLA4 has seven neighbors and six of them were CKGs. Such a close connection with CKGs suggested these genes as potential drug targets for BLCA treatment.

### Sensitive Cancer-keeping genes as potential therapeutic targets for BLCA

We subjected six sCKGs to *in vitro* and one sCKGs *in vivo* analyses to assess their impact on BLCA cell proliferation and migration, thus evaluating and confirming their function in maintaining cancerous cell states. We chose a) four sCKGs (RPS6KA3, N-cadherin (CDH2), caspase-1, and FN1) that are not cancer driver genes for BLCA or any other malignancies to assess their potential function in BLCA, and b) two sCKGs (FGFR3 and EP300) that are known cancer driver genes for BLCA^46^ for comparison and validation (Figure 3D, gene names marked in red). We adopted small-interferencing RNAs (siRNAs) to knock down these six sCKGs in two BLCA cell lines (T24 and UMUC3) and one sCKG (RPS6KA3) in a mouse model of BLCA.

The siRNA knockdown experiments were repeated three times for every gene and every cell line, and the experiments were also repeated on ten mice. For each experiment, the expression of the sCKG was measured by quantitative PCR. The results of the repeated experiments were consistent (Figure 4A). The absence of RPS6KA3, FGFR3, and CDH2 significantly decreased the proliferation of the two BLCA cells (Figures 4A and 4B). In contrast, the knockdown of EP300 and FN1 significantly promoted the proliferation of UMUC3 cells (Figure 4B) but had little effect on the proliferation of T24 cells (Supplemental eFigures 11A and 11B). Interestingly, the loss of caspase-1, which was thought to play an important role in inflammation and tumor microenvironment, promoted the proliferation and survival of BLCA cells (Figures 4A and 4B). Moreover, the deletion of FGFR3, RPS6KA3, EP300, FN1, and CDH2, as expected, significantly reduced the migration ability of bladder tumor cells (Figures 4C and 4D). In contrast, the absence of caspase-1 promoted the migration of BLCA cells (Figures 4C and 4D).

**Figure 4.**
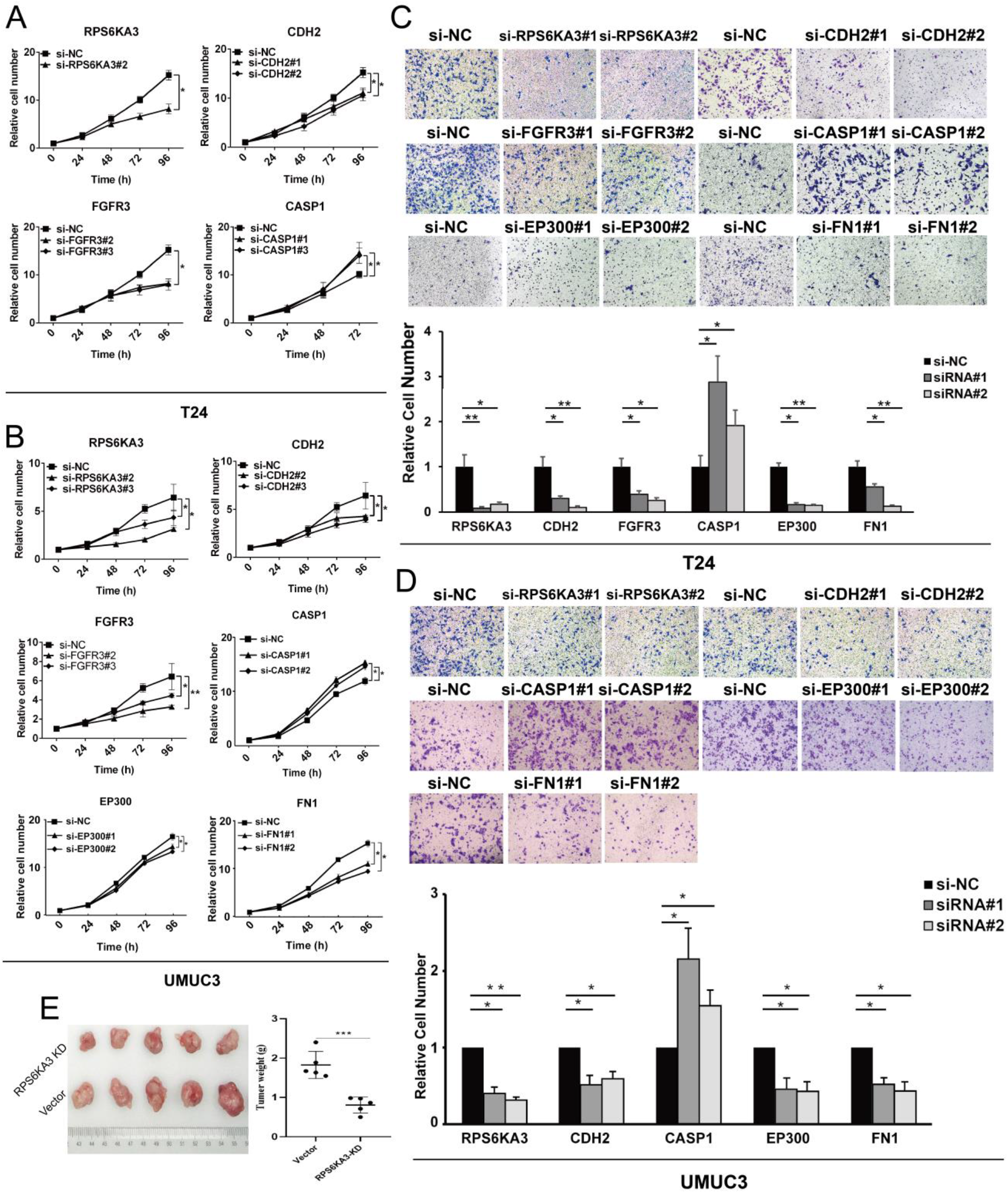
Experimental validation of representative sensitive CKGs (sCKGs) in bladder cancer. **A-B)** CCK-8 assay and corresponding quantitative detection of the effect of siRNA-mediated knockdown sCKGs on the proliferation of T24 (A) and UMUC3 (B) BLCA cells. **C-D)** Transwell migration assay (upper) and corresponding quantitative detection (down) of the effect of siRNA-mediated knockdown sCKGs on the migration ability of T24 (C) and UMUC3 (D) cells. **E)** RPS6KA3 promoted tumor proliferation *in vivo*. BALB/c nude mice were injected with UMUC3 cells which were stably transfected with RPS6KA3 knockdown plasmids or control vector in the xenotransplantation model. After 4 weeks, the tumor size (left) and weight (right) in the RPS6KA3 knockdown group were significantly lower than those of the vector group. Data represent mean±s.d. from three replicate cultures. P-values were computed using the one-sided t-test.

Furthermore, we used a nude mouse model with transplanted BLCA tumors to examine the function of a typical sCKG, RPS6KA3, which has Serine/Threonine kinase activities and plays an important role in the proliferation of some solid tumors^48, 50^. Consistent with the *in vitro* results, the knockdown of RPS6KA3 in the mouse model significantly inhibited the growth of BLCA xenografts in the mice (Figure 4E). Putting together, all the *in vitro* and *in vivo* experimental results confirmed the functional roles of some of the sCKGs for maintaining the validity of cancer cells and validated the feasibility of our novel network control hub-based method.

## DISCUSSION

One of our major contributions was a novel perspective on how to view and treat cancer. It is fundamentally different from the prevalent concept of cancer-driver genes (CDGs) that may transform normal cells into abnormal ones with uncontrolled cell growth. We introduced a new notion of cancer-keeping genes (CKGs) defined on cancerous cells which when intervened may terminate or prevent abnormal cell differentiation and proliferation. To utilize the new concept of CKGs, we resorted to and extended the well-developed theory of network structural controllability^11^, which has been adopted to identify CDGs and cancer therapeutic targets^33, 51^. We extended one control scheme in structural controllability to define control hubs over all control schemes. We adopted control hubs as candidates of CKGs defined for cancerous or abnormal cells.

There are typically much fewer control hubs (or CKGs) than driver nodes in a gene regulatory network. For example, BLCA_GRN has 5,052 driver genes out of a total of 7,030 genes but only 660 CKGs can be further reduced to 173 sCKGs. Most CKGs for BLCA are involved in important tumor signaling pathways. Remarkably, the genes in the cell-cycle and p53 signaling pathways as reported by the TCGA project^38^ are all CKGs (Figure 2F). Dysregulation of cell-cycle control is a hallmark of cancer^52^. Many CKGs in the cell-cycle pathway, including CDKN1A^53^, CDK2^54^, and E2F1^55^, have been considered potential targets of anticancer drugs. The p53 pathway is tightly regulated in BLCA^56^ and the genetic variations of the genes in the pathway have been an attractive topic for BLCA treatment^57, 58^. CDKN1A and TP53 in the cell-cycle and p53 pathways, respectively, have been taken as therapeutic targets for BLCA^59^. The enrichment of the CKGs in TGFß and receptor-tyrosine kinase (RTK)/RAS/MAP-Kinase signaling pathways is also thought-provoking. The two pathways are known to be important for BLCA. TGFß can act as a critical tumor suppressor^60^ and the dysregulation of the TGFß pathway may increase the risk of BLCA. Sixteen CKGs are also involved in other cancer-related signaling pathways, including PI3K, Myc, Wnt, Notch, Hippo, and Nrf2 signaling pathways (Supplemental eTable 4).

We like to highlight that CKGs are network-structure-based so they are fundamentally different from mutation-based CDGs. Most known CDGs have high mutation rates and/or high connectivity in gene regulatory networks. In contrast, CKGs, particularly sCKGs, have average mutation rates and thus are unlikely to be detected by the existing mutation-based methods. On the other hand, most CKGs are directly connected to known CDGs and drug targets in BLCA_GRN, forming gene pairs that are essential for the maintenance and function of the network. Many CKGs are also located upstream in tumor-related signaling pathways and thus can control CDGs. Therefore, CKGs provide alternatives to CDGs, and many CKGs can be viewed or adopted as latent, master regulators of CDGs. Indeed, manipulating CKGs can affect cancer cell proliferation and migration and suppress tumor growth, as we showed in our experiments using two cell lines and a mouse model of BLCA (Figure 4). The six sCKGs (RPS6KA3, FGFR3, N-cadherin (CDH2), EP300, caspase-1, and FN1) that we experimentally analyzed are neither known CDGs of BLCA nor drug targets for BLCA and have not been well studied for bladder cancer. Combined, our analytic and experimental results strongly suggest CKGs be a class of novel regulatory elements that, when perturbed, can affect the states of the underlying cells. In particular, the six experimentally analyzed sCKGs are novel putative drug targets for BLCA treatment.

Sensitive control hubs or sCKGs have different characteristics from the conventional network hub nodes defined by various structure-based methods, such as the nine popular methods that we analyzed (eFigure 7). Sensitive control hubs are not typical network hubs and have lower centrality, therefore, most of them would not be detected by the existing hub-based methods. The sensitive control hubs are essential to network control and any variation to them may change overall network states. Meanwhile, the lower centrality of sensitive control hubs makes variations to them have less impact on network (or cell) stability than the other methods. Therefore, they are excellent candidates for therapeutic targets.

The extension to network structural controllability as done in our work is thought-provoking and is expected to inspire future work to extend the theory of structural controllability in multiple dimensions for the urgent demands for innovative methods for analyzing large quantities of biological data. Our novel control-hub-based analytic approach can be directly applied to various types of cancer and be readily extended to other complex diseases, such as neurodegenerative disorders and metabolic diseases.

## MATERIALS and METHODS

### Data of genes, functions, and pathways used

Five well-curated disease- and cancer-related pathway databases were used to construct a bladder cancer gene regulatory network (BLCA_GRN). In particular, the ten most important and common types of interactions (Supplemental eTable 1) were used to link genes. The five databases included the National Cancer Institute (NCI) Nature Pathway Interaction Database^24^, PhosphoSite Kinase-substrate information^25^, HumanCyc^26^ (https://humancyc.org), the Reactome^27^ (https://www.reactome.org) and PANTHER Pathways^28^ (http://www.pantherdb.org). These database projects were in part motivated by disease studies, so they contained information on mutated genes and disease and cancer-related pathways. For example, Reactome consists of the KEGG databases that include a disease database and a drug database. Phosphosite contains data on missense mutations from UniProtKB, TCGA, and many other sources, which were collected from more than 2,000 diseases and syndromes, and has data on polymorphisms associated with hundreds of cancers.

### Control scheme and control hubs of cancer gene regulatory network

A network (or cell) can be structurally controlled by exerting external signals on control nodes (or driver genes) so that it can be driven from any initial state to the desired state^10^. A minimal set of control nodes can be found by the maximum matching of the nodes in the network. A maximum matching of the network is a maximal set of edges that share no node in common. The nodes without matching edges directly connected are control nodes. From the control nodes (or head nodes), moving along matching edges form control paths. All the control nodes and control paths form a control scheme of the network^31^.

Since maximum matching is not unique^21, 22^, a network often has multiple control schemes. A node may occupy distinct positions in control paths of different control schemes. For example, a control node may become a node in the middle of a control path of another control scheme. A node is a *control hub* if it resides in the middle of a control path in *every* control scheme of the network^31^. All control hubs can be identified as follows. Firstly, for a directed bipartite network *B(V_in_, V_out_, E)*, all possible driver nodes can be marked in the process of finding all augmented paths, denoted as *H*. The method for computing all possible driver nodes has been proved in detail in our previous work^31^. Next, find all possible driver nodes of transpose network *B(V_out_, V_in_, E)*, denoted as *T*. All control hubs of a network can be identified by excluding all driver nodes *H* and all tail nodes *T* in polynomial time without computing all control schemes^31^.

### Identification of sensitive control hub nodes or sensitive cancer-keeping genes

While control hubs are vulnerable spots for the structural controllability of a network, they may have different resilience to external stimuli. A control hub is referred to as being *sensitive* if it is no longer a control hub after removing any edge from the network. All sensitive control hubs of a network can be identified by removing one edge at a time and examining whether the set of control hubs of every control scheme remained intact. The complexity of this algorithm is O(*n*^0.5^*m*^2^) on a network with *n* nodes and *m* links in the worst.

### Functional enrichment analysis of cancer-keeping genes

A function enrichment analysis was adopted to assess the biological functions of CKGs using the information in 32 gene function datasets (Supplemental eTable 3). The enrichment for a particular function was reflected by those CKGs that were annotated with that function. To evaluate the statistical significance of the enrichment, a z-test was performed as follows. A sample of *N* genes was randomly chosen from the genes in BLCA_GRN, where *N* is the number of CKGs. The functional enrichment of this random sample was the overlapping of the random set with the given functional dataset. Random samplings were chosen 1,000 times, and their enrichments constituted an empirical enrichment distribution of CKGs. A *z-test*, using 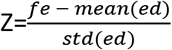, was adopted to measure the difference between the function enrichment of CKGs (*fe*) and the empirical distribution (*ed*), along with the statistical significance of the two-tailed p-value.

### Map sensitive CKGs with cancer driver genes and cancer therapeutic targets

The CKGs were further analyzed using several datasets including that of CDGs, cancer therapeutic targets, and immune genes (eTable 3). The CDG dataset^29^ includes 739 genes from 9,423 tumor exome sequencing data from TCGA. Cancer therapeutic target dataset^45^ includes 628 genes obtained from genome-scale CRISPR-Cas9 screening of 324 human cancer cell lines from 30 cancer types. Immune-related genes include 358 genes obtained by coreCTL immune-related gene dataset^30^, CAR immune-related gene datasets^31^, and immune checkpoints^36, 37^. We took the overlapping genes among the three datasets with the CKGs for further analysis.

### Survival analysis

Survival analysis was performed on 35 sCKGs using the clinical data of 411 samples of Bladder Urothelial Carcinoma (TCGA PanCancer analysis) from the cbioportal website (http://www.cbioportal.org). We counted the number of mutated genes in 35 sCKG across all 411 samples and selected 4 subsets based on the number of simultaneously mutated genes. Whether a gene was considered mutated depends on the common alterations (somatic mutation, gene fusion, copy number amplification, or homozygous deletion) it contained. These subsets respectively had no mutated genes, at least one, two, and three mutated genes in the 35 sCKGs. The survival analysis was performed using the Kaplan-Meier estimator^61^ from the cbioportal website (http://www.cbioportal.org).

### Cell culture

Human bladder cancer-derived UMUC3 and T24 cells were purchased from the Chinese Academy of Sciences (Shanghai, China). UMUC3 and T24 cells were cultured in DMEM (11995-065, Gibco, USA) and DMEM-F12 (10565-018, Gibco), respectively. These mediums were supplemented with 10% heat-inactivated fetal bovine serum (FBS; 1750114, Gibco). Cells were maintained at 37°C in an incubator containing 5% CO_2_.

### Transient siRNA-mediated gene knockdown

The negative control (siRNA-NC) and siRNAs targeting the CKGs to be tested were purchased from Shanghai Genechem Co., Ltd. (China). For transient transfection, a designated siRNA was introduced into cells in the presence of Lipofectamine 3000 transfection reagent (Invitrogen, Carlsbad, CA, US) according to the manufacturer’s instructions. The siRNA was diluted to the final concentration of 20 μM following the manufacturer’s instructions. Cancer cells were transfected using Lipofectamine TM RNAiMAX (Invitrogen, Carlsbad, CA, US) and opti-MEM (Gibco). Twenty-four hours after transfection, cells were collected and subjected to subsequent experiments. The sequences of the siRNAs tested are listed in Supplemental eTable 10-11. Three siRNAs were designed to target different regions of each target gene tested, and the knockdown efficiency was tested. The siRNA with a significant knockdown effect was selected for subsequent functional verification and the rest siRNAs were discarded.

### Cell Counting Kit 8 (CCK-8) assay

The effect of knocking down candidate CKGs on cell proliferation was examined by CCK-8 assays. Twenty-four hours after transfection, cells were collected and maintained in 96-well plates (1,500 cells/well). Four hours after incubation, 10 μl of CCK-8 reagent (Vazyme, Nanjing, China) was added to each well and the reaction mixtures were incubated for 2 h at 37 °C under a 5% CO_2_ atmosphere. CCK-8 assay was performed at the same time every day for 4 consecutive days, and a total of 96h of cell activity was recorded. The absorbance (OD) value at 450 nm wavelength was then measured using a Microplate reader (Bio-Rad, CA).

### Transwell assay

Transwell assays were employed to determine the effect of candidate cancer-keeping genes on the migration and invasion abilities of cancer cells. Twenty-four hours after transfection, cells were collected and subjected to transwell assay. In brief, the upper compartment of the filters (Corning, NY, USA) was coated with (invasion) or without (migration) 55 μl of the basal membrane matrix (1:7 dilution, Corning). Cells were diluted with serum-free medium to 1× 10^5^ cells per ml. Cell suspension (200 μl) was added to the upper compartment, and 600 μl of RPMI-1640 containing 10% FBS was added to the lower compartment. Twenty-four hours after incubation at 37°C under 5% CO_2_, filters were fixed with 4% formaldehyde for 30 min, then stained with 0.1% crystal violet at room temperature for 30 min. Filters were then rinsed 3 times with PBS, and the unmigrated cells were removed with cotton swabs. Finally, cells were photographed and counted in a 5-independent microscopic field.

### Xenograft mouse model

UMUC3 cells expressing control shRNA or PRS6KA3 shRNA (2×10^6^) were subcutaneously injected into the dorsal flank of 4-week-old male athymic nude mice (*n*=5 mice per group, Shanghai SLAC Laboratory AnimalCo. Ltd.). Mice were sacrificed after 3 weeks, and tumors were excised and weighed. Mice were used in the experiment at random. During testing the tumors’ weight, the researchers were blinded to the information and shape of tumor tissue masses. Studies on animals were conducted with approval from the Animal Research Ethics Committee of China Medical University.

## Data and software availability

The datasets used in the study are in Supplement File 1 and Supplement File 2. The datasets and software of our method for finding control hub nodes and cancer-keeping genes are freely available in the public software repository GitHub at https://github.com/network-control-lab/control-hubs.

## Author contributions

Xizhe Zhang conceived and designed the project as well as supervised the work except for the biological experiments. Xizhe Zhang also designed the algorithm, analyzed its correctness and complexity, performed data analysis, and draft the manuscript. Weixiong Zhang conceived the main conceptual ideas and draft the manuscript. Chunyu Pan and Xinru Wei implemented the algorithm and collected and analyzed the data. Yuyan Zhu designed and supervised the *in vivo* and *in vitro* experiments and performed data analysis and integration and related manuscript writing. Yu Meng was involved in the design of *in vivo* experiments and related data analysis. Shang Liu, Jun An, and Jieping Yang carried out the *in vitro* and *in vivo* experiments and data collection. Baojun Wei and Wenjun Hao analyzed the data and results from the *in vitro* and *in vivo* experiments.

## Acknowledgment

This work was supported in part by the National Natural Science Foundation of China (grant numbers 62176129 and 81672523), the United States National Institutes of Health (grant number R01-GM100364), and the Hong Kong Global STEM Professorship Scheme.

