## Supplemental material for "Cancer-keeping genes as therapeutic targets"

### This file includes:

eMethods 1. Structural controllability of directed network
eMethods 2. Identifying control hubs of complex networks

eFigure 1. An example of control paths of a network and its transpose network
eFigure 2. Enrichment analysis of essential genes and conserved genes
eFigure 3. Enrichment analysis of signaling proteins and protein abundance
eFigure 4. Enrichment analysis of protein PTMs
eFigure 5. Enrichment analysis of disease genes, human viruses and drug targets
eFigure 6. Enrichment analysis of and immune genes
eFigure 7. Enrichment analysis of control hub and other network hub identification methods
eFigure 8. The distribution of neighbors of 173 sCKGs
eFigure 9. The distribution of 173 sCKGs in different datasets
eFigure 10. The network of 35 sCKGs and their neighbor genes
eFigure 11. Experimental validation of representative sCKGs in cervical cancer and head and neck cancer cells

eTable 1. The types of control-related interactions
eTable 2. Seed nodes of BLCA gene regulatory network
eTable 3. Datasets for functional enrichment analysis
eTable 4. The enrichment of the CKGs in ten important cancer signaling pathways
eTable 5. The sensitive edges and their corresponding confidence scores
eTable 6. Neighbor Cancer driver of the 35 sCKGs
eTable 7. Neighbor Drug Targets of the 35 sCKGs
eTable 8. Neighbor Immune gene of the 35 sCKGs
eTable 9. Six drug targets and their neighbors of BLCA
eTable 10. Sequences of oligonucleotide primers used for qPCR in the study
eTable 11. Sequences of oligonucleotide siRNA or shRNA used in the study

**Other Supplementary Materials for this manuscript include:**

Tables File 1
Tables File 2

**eMethods 1. Structural controllability of directed network**

According to control theory, a dynamic networked system can be driven from any initial state to any final state in finite time if it was inputted with a suitable choice of inputs and external control signals[1]. To identify the controllability and find suitable inputs for a network, structural controllability was introduced by Lin[2] and developed by Liu et.al[1], which maps the problem of finding the minimum inputs or driver nodes into identifying a maximum matching of the network. Although most biological networks are inherently nonlinear, the experimentally obtained network, however, these networks can be assumed to capture linear effects around homeostasis. Therefore, we used the concept of local structural controllability[1] and the analysis tools based on linear controllability to investigate the controllability of the cancer gene regulatory network.

Consider a directed network $G(V, E)$, the network can be modeled as a linear time-invariant system, whose states are determined by the following equations (1):

$$\frac{dx(t)}{dt} = Ax(t) + Bu(t) \tag{1}$$

where the state $\mathbf{x}(t)=(x_1(t),\ldots, x_N(t))^T$ denotes the state value of all nodes at time $t$; $A$ is the transpose of the adjacency matrix of G; $\boldsymbol{u}(t)=(u_1(t),\ldots,u_M(t))^T$ is the input signal; $B$ is the input matrix that defines how control signals are inputted into the network. The nodes receiving external signals are called **driver nodes**. The minimum set of driver nodes to fully control the network is called **M**inimum **D**river nodes **S**et (*MDS*). Based on the minimum input theorem present by Liu et.al[1], the *MDS* can be obtained by computing any maximum matching of an equivalent undirected bipartite graph $B(V_{in}, V_{out}, E)$[3]. The set of unmatched nodes of $V_{in}$ are the *MDS*. While solving the maximum matching process as mentioned above, the network is transformed into a bipartite graph by splitting the node set $V$ into two node sets $V_{in}$ and $V_{out}$ and the maximum matching is obtained by finding the augmented path.

Next, we will introduce some concepts in graph theory. A set of edges in $B$ ($V_1$, $V_2$, $E$) is called a **matching** $M$ if no two edges in $M$ have a node in common[4]. A node $v_i$ is said to be **matched** by $M$ if there is an edge of $M$ linked to $v_i$; otherwise, $v_i$ is unmatched[3]. The maximum matching is a matching with the maximum number of edges of the graph. A path $P$ is said to be $M$-alternating if the edges of $P$ are alternately in and not in $M$. An $M$-alternating path $P$ that begins and ends at the unmatched nodes is called an $M$-augmenting path. The maximum matching is a matching with the maximum number of edges of the graph.

**eMethods 2. Identifying control hubs of complex networks**

Based on structural controllability theory, for a directed network G(V, E), the matching edges of a maximum matching form the cactus structures in the network, which are the basic control structure of the network. Therefore, the matched edges form a set of edge-independence paths in the directed network $G(V, E)$, we call these paths as ***control paths***[5]. The control paths start with driver nodes and end with tail nodes. The driver nodes (unmatched nodes) and the corresponding control path in the network are called a ***control scheme***.

Considering a node in the network, based on its position on the control path, we can classify it into three types. A node is called a ***head node*** if it is at the beginning of a control path of at least one control scheme. Similarly, a node is a ***tail node*** if it is at the end of a control path of at least one control scheme. A node is a ***middle node*** if it is in the middle of a control path of at least one control scheme. However, the maximum matching is not unique[6] for most networks and many control schemes exist in a network. A node may change its type in different control schemes, i.e., a head node in one control scheme may become a middle node in another control scheme. Therefore, we consider a special type of node, which always lies in the middle of a control path in all control schemes. We call these nodes ***control hub nodes***.

We were interested in identifying control hub nodes, which are middle nodes in all control schemes. If a node is not a head node or tail node in all control schemes, it must be a control hub. Based on the above definition, we need to first find out the all head and tail nodes in all control schemes, then the rest nodes are the control hub nodes. To find head or tail nodes, we have the following claims:

**Property 1**. For a given network G and all its control schemes, the head nodes are the union of all driver nodes of all control schemes.

**Proof**: Based on the definition of the driver nodes, the external control signals are inputted into the network from the driver nodes. Therefore, the driver nodes lie at the beginning of the control paths of a control scheme. Therefore, the head nodes are the union of all the driver nodes of all control schemes of a network.

Consider a network G, we say a network G' is the transpose network of G, if it has the same nodes set but the direction of all edges is reversed compared to the corresponding edges in G. To identify tail nodes, we have the following property:

**Property 2**. For a given network *G* and its transpose network *G*', the head nodes of G' are the tail nodes of G.

**Proof:** For the directed network G(V, E), let B ($V_{in}$, $V_{out}$, $E$) is the corresponding undirected bipartite graph. Consider the transpose network *G*'(V, E), the only difference between G and G' is the direction of all edges are reversed (see eFigure for an example ). Therefore, the undirected bipartite graph of G' must be B' ($V_{out}$, $V_{in}$, $E$). Because B' and B have the same edges set, therefore, a maximum matching of $B$ ($V_{in}$, $V_{out}$, E) must be a maximum matching of B' (V $V_{out}$, $V_{in}$, E). Therefore, the control paths of all control schemes of G are the same as that of G' except they have reversed direction, which make the head nodes of G' must be the tail nodes of G. The proof is completed.

**Theorem 1:** For a network G(V, E), the set of control hubs C=V-H-T, where H is the set of the head nodes and T is the set of the tail nodes of the network.

**Proof**: It is trivial based on the definition of control hubs.

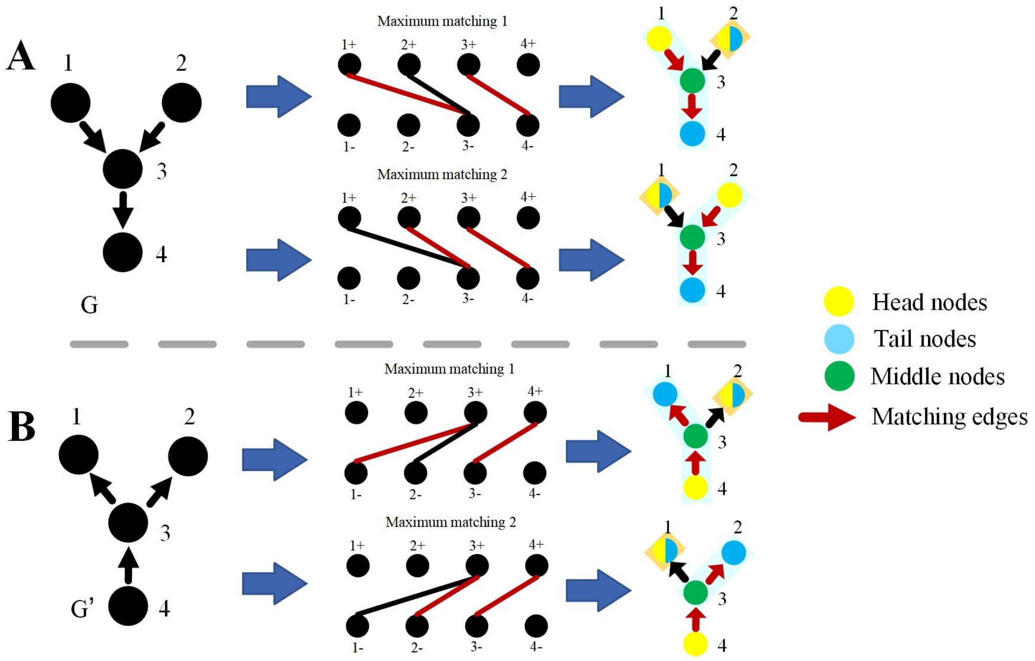


**eFigure 1. Network G and its transpose network G’.** For a given network G and its transpose network G’, the head nodes of G’ are the tail nodes of G. The difference between their bipartite graphs is the exchange of node sets, but they will obtain the same maximum matching and control paths.

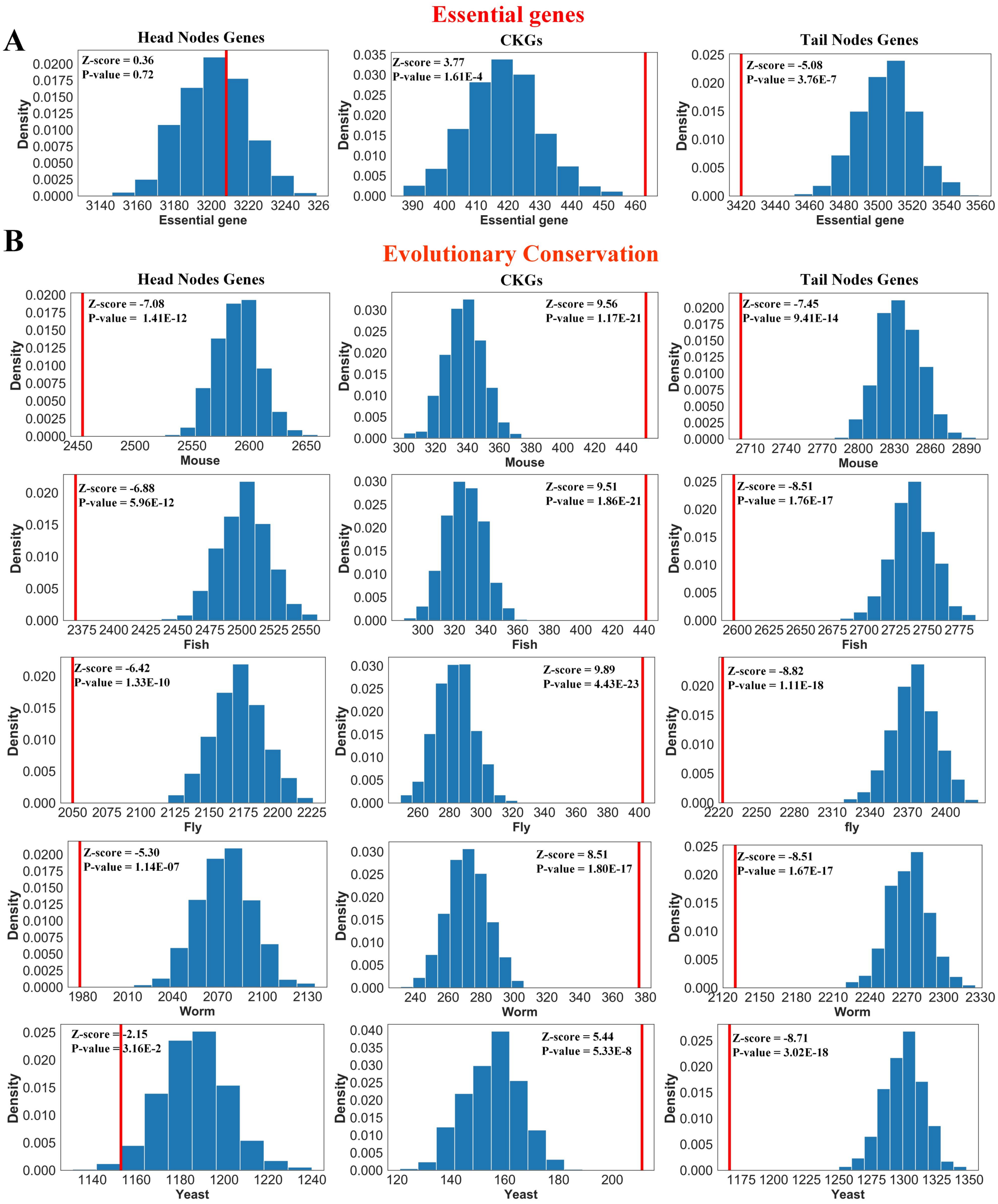


**eFigure 2.** Enrichment analysis of essential genes and conserved genes in the three types of nodes. **A)** Enrichment analysis of essential genes. Numbers of essential genes overlapping with head, mid and tail nodes are shown in red lines, and their respective size-controlled random set distributions are in blue bars. Essential

genes were significantly enriched in the mid-type nodes by comparing z-score; **B)** Enrichment analysis of evolutionary conservation genes. The numbers of genes conserved in mice, fish, flies, worms, and yeast are shown in the red lines, and their respective size-controlled random set distributions are shown in blue bars. Conserved genes were also significantly enriched in the mid-type nodes by comparing the z-score.

**A**

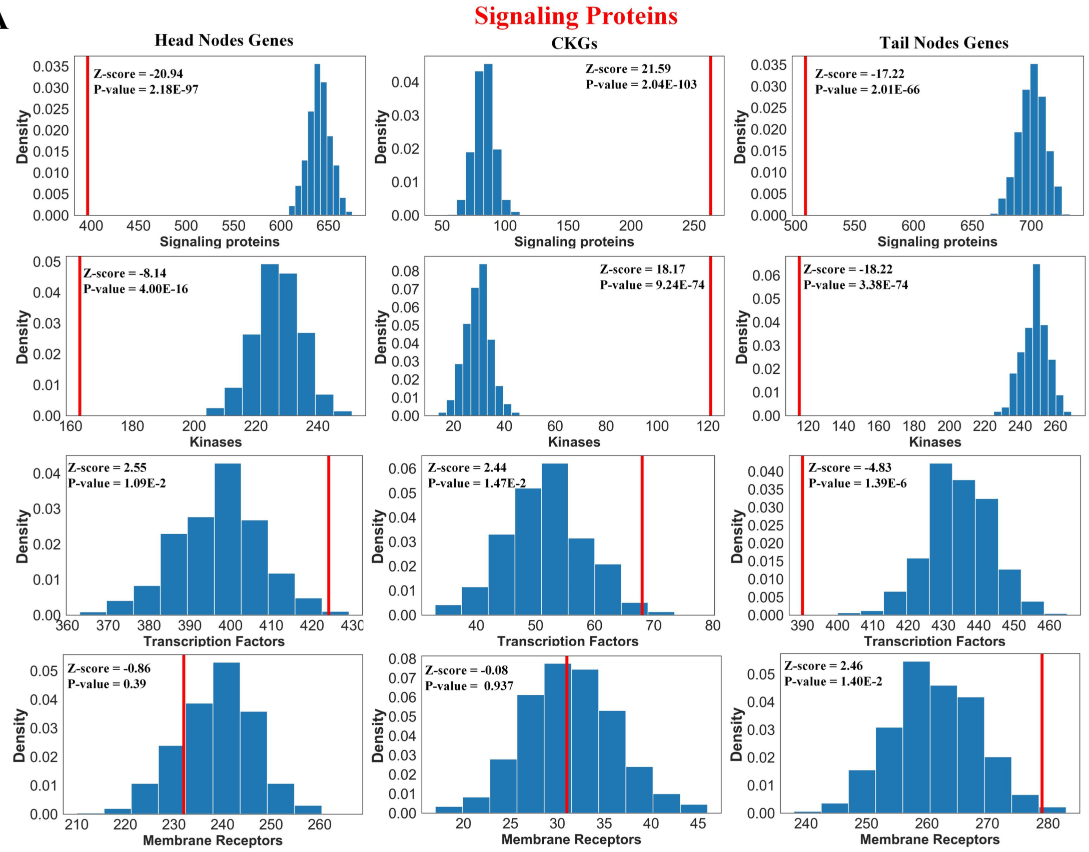
Signaling Proteins
Head Nodes Genes
CKGs
Tail Nodes Genes
Z-score = -20.94
P-value = 2.18E-97
Z-score = 21.59
P-value = 2.04E-103
Z-score = -17.22
P-value = 2.01E-66
Signaling proteins
Z-score = -8.14
P-value = 4.00E-16
Z-score = 18.17
P-value = 9.24E-74
Z-score = -18.22
P-value = 3.38E-74
Kinases
Z-score = 2.55
P-value = 1.09E-2
Z-score = 2.44
P-value = 1.47E-2
Z-score = -4.83
P-value = 1.39E-6
Transcription Factors
Z-score = -0.86
P-value = 0.39
Z-score = -0.08
P-value = 0.937
Z-score = 2.46
P-value = 1.40E-2
Membrane Receptors
Density

**B**

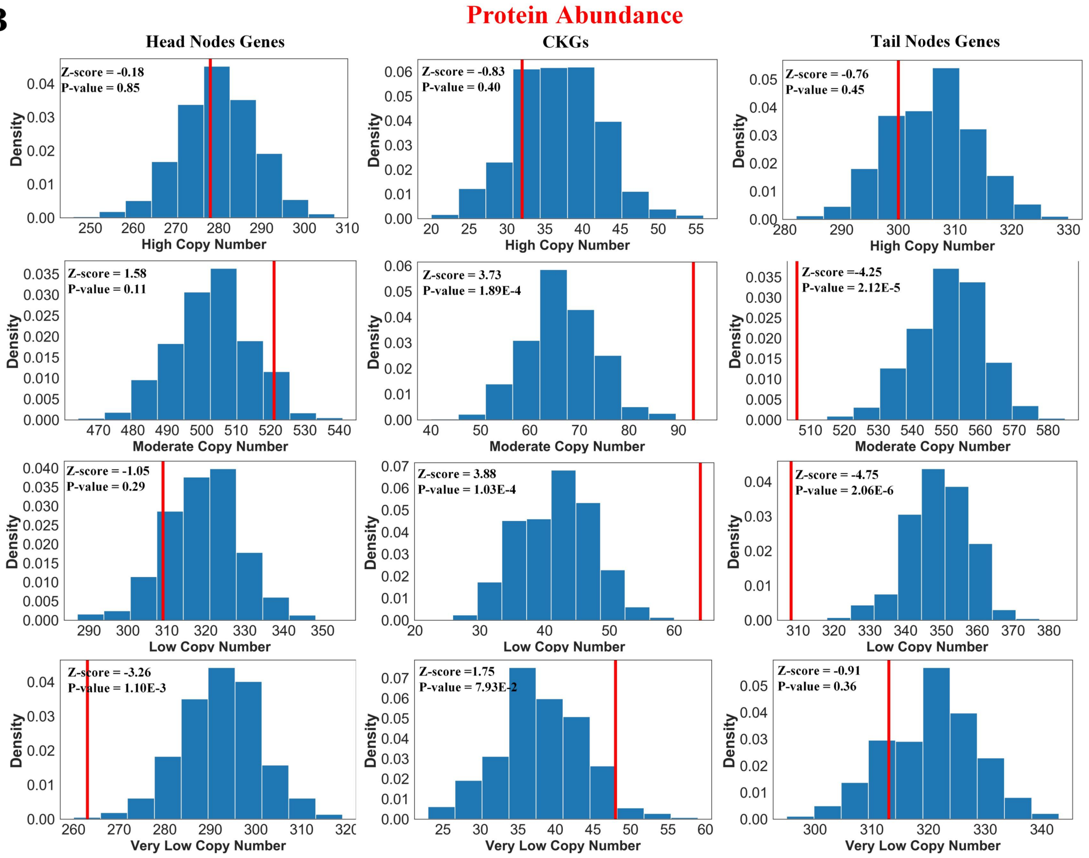
Protein Abundance
Head Nodes Genes
CKGs
Tail Nodes Genes
Z-score = -0.18
P-value = 0.85
Z-score = -0.83
P-value = 0.40
Z-score = -0.76
P-value = 0.45
High Copy Number
Z-score = 1.58
P-value = 0.11
Z-score = 3.73
P-value = 1.89E-4
Z-score = -4.25
P-value = 2.12E-5
Moderate Copy Number
Z-score = -1.05
P-value = 0.29
Z-score = 3.88
P-value = 1.03E-4
Z-score = -4.75
P-value = 2.06E-6
Low Copy Number
Z-score = -3.26
P-value = 1.10E-3
Z-score = 1.75
P-value = 7.93E-2
Z-score = -0.91
P-value = 0.36
Very Low Copy Number
Density

**eFigure 3.** Enrichment analysis of signaling proteins and protein abundance in the three types of nodes. **A)** Enrichment analysis of signaling proteins. Numbers of nodes overlapping with signaling proteins, receptors, protein kinases, and transcription factors are shown in the red lines, and their respective size-controlled random set distributions in blue bars; **B)** Enrichment analysis of protein abundance. The numbers of nodes overlapping with high copy numbers, moderate copy numbers, low copy numbers, and very low copy numbers are shown in red arrows and their respective size-controlled random set distributions are in blue bars.

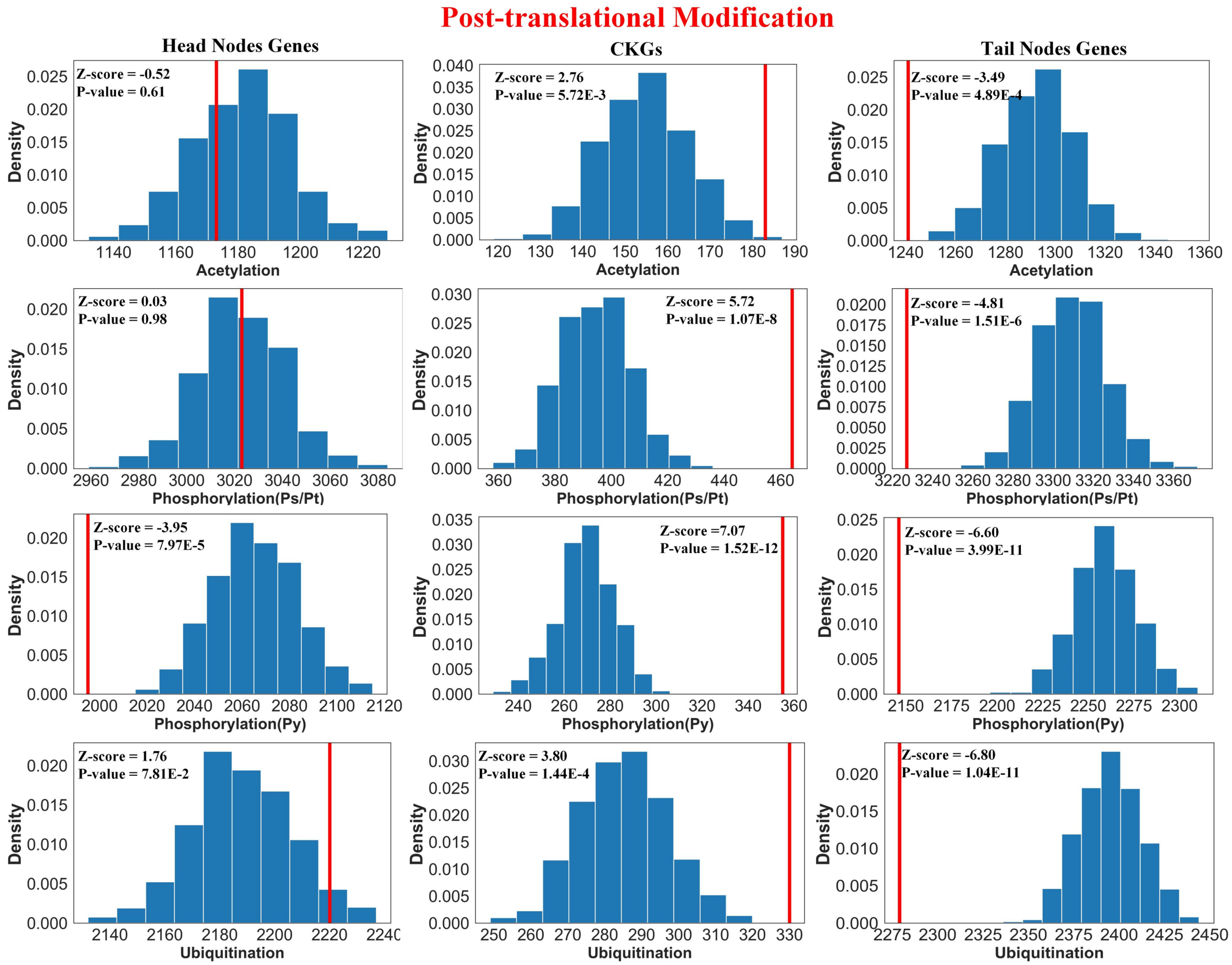


**eFigure 4.** Enrichment analysis of protein PTMs in the three types of nodes**.** Numbers of nodes overlapping with Acetylation, Tyrosine Phosphorylation, Serine/Threonine Phosphorylation, and Ubiquitination datasets are shown in red lines, and their respective size-controlled random set distributions in blue bars.

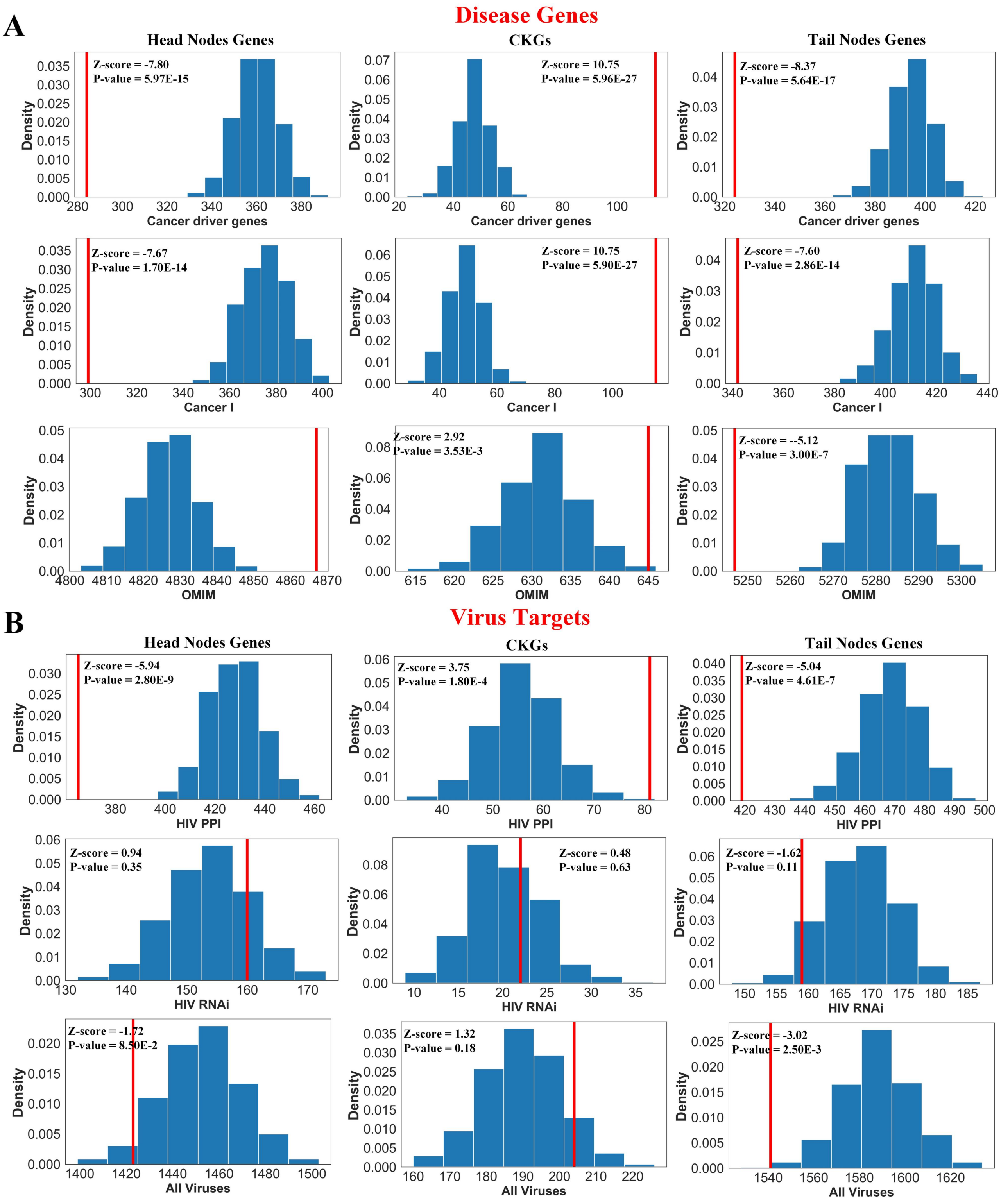
A
Disease Genes
Head Nodes Genes
CKGs
Tail Nodes Genes
Z-score = -7.80
P-value = 5.97E-15
Z-score = 10.75
P-value = 5.96E-27
Z-score = -8.37
P-value = 5.64E-17
Cancer driver genes
Z-score = -7.67
P-value = 1.70E-14
Z-score = 10.75
P-value = 5.90E-27
Z-score = -7.60
P-value = 2.86E-14
Cancer I
Z-score = 2.92
P-value = 3.53E-3
Z-score = --5.12
P-value = 3.00E-7
OMIM
Density
B
Virus Targets
Head Nodes Genes
CKGs
Tail Nodes Genes
Z-score = -5.94
P-value = 2.80E-9
Z-score = 3.75
P-value = 1.80E-4
Z-score = -5.04
P-value = 4.61E-7
HIV PPI
Z-score = 0.94
P-value = 0.35
Z-score = 0.48
P-value = 0.63
Z-score = -1.62
P-value = 0.11
HIV RNAi
Z-score = -1.72
P-value = 8.50E-2
Z-score = 1.32
P-value = 0.18
Z-score = -3.02
P-value = 2.50E-3
All Viruses

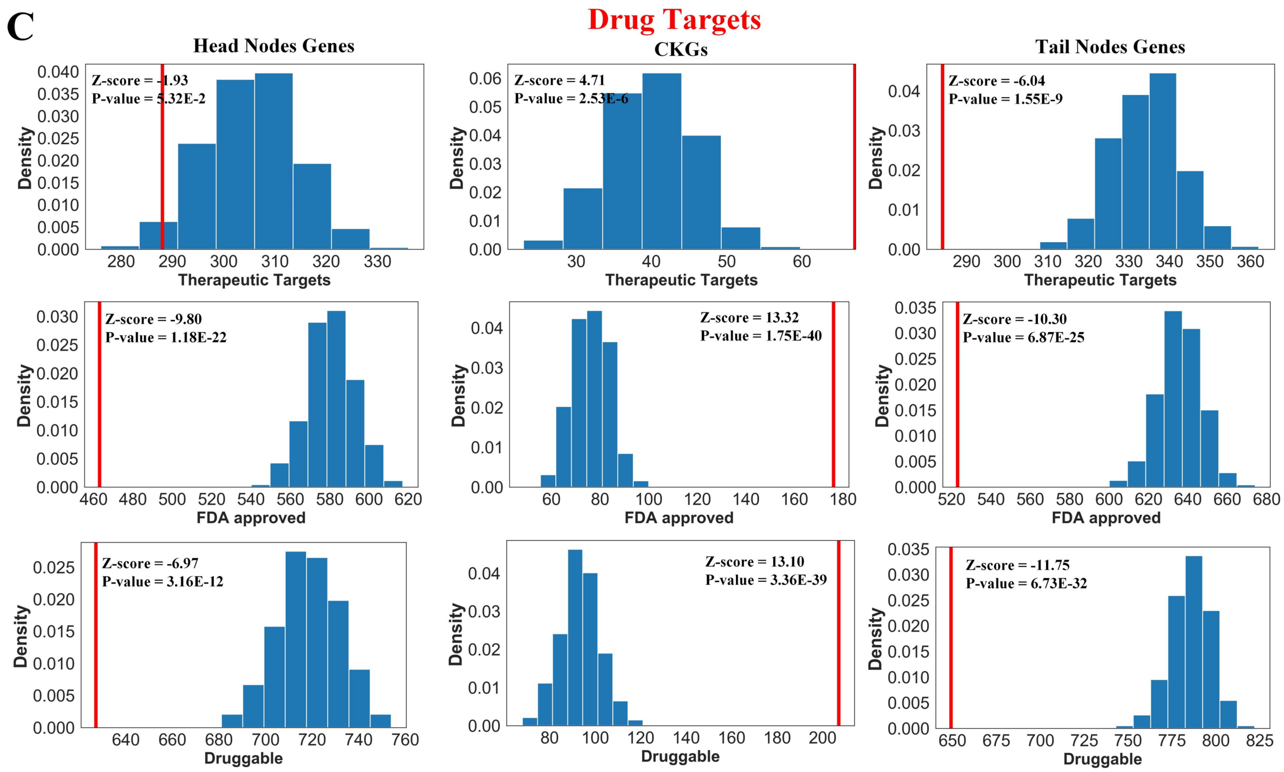


**eFigure 5.** Enrichment analysis of disease genes, human viruses, and drug targets in the three types**. A)** Enrichment analysis of Disease Genes. Numbers of nodes overlapping with Cancer driver genes, Cancer I and OMIM are shown in the red lines and their respective size-controlled random set distributions. **B)** Enrichment analysis of human viruses. Numbers of nodes overlapping with HIV and Viruses are shown in the red lines and their respective size-controlled random set distributions. **C)** Enrichment analysis of Drug Targets. Numbers of nodes overlapping with Therapeutic Targets, FDA-approved drug targets, and Druggable genes are shown in the red lines and their respective size-controlled random set distributions.

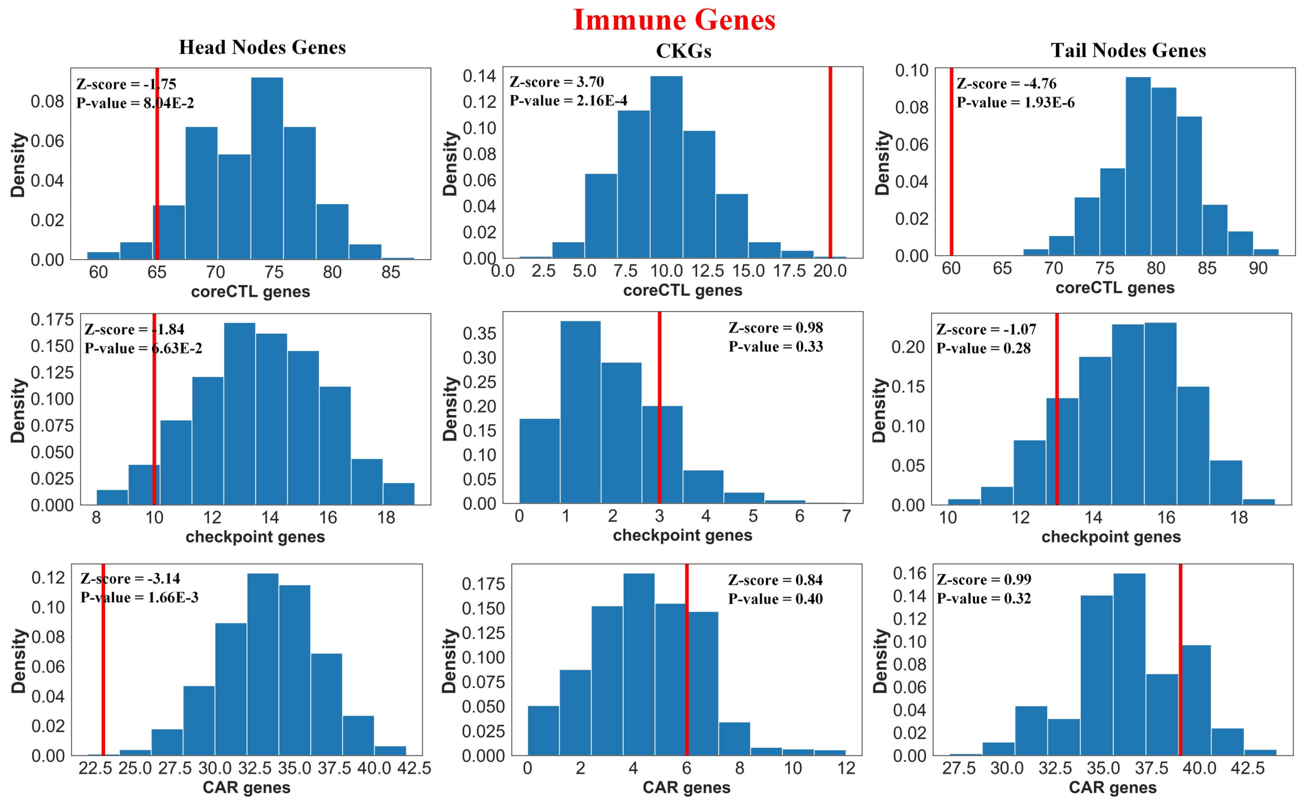


**eFigure 6.** Enrichment analysis of immune genes in the three types. Numbers of nodes overlapping with cancer innate immune escape genes (coreCTL genes), CAR therapy targets, and immune checkpoints are shown in red line and their respective size-controlled random set distributions are in blue bars.

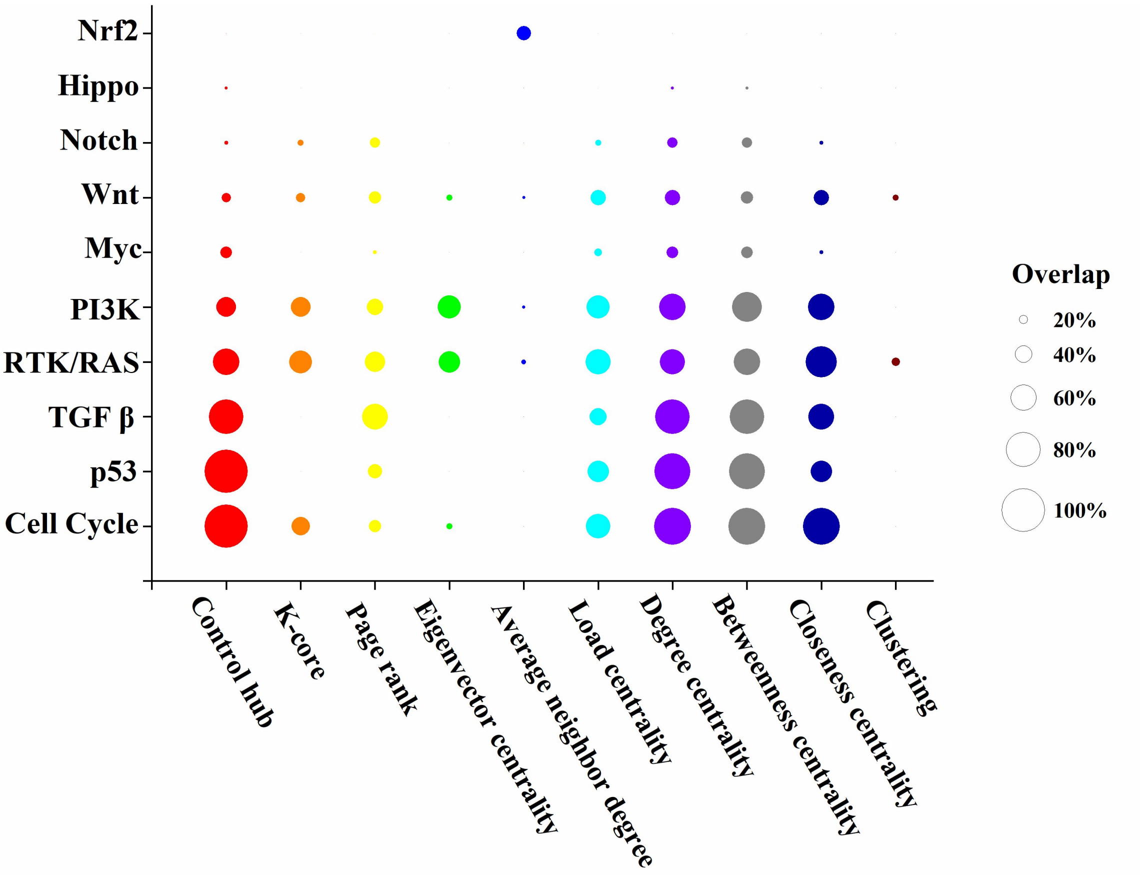


**eFigure 7.** Enrichment analysis of control hub and nine popular network hub identification methods. The horizontal coordinate lists the different network hub identification methods and the vertical coordinate shows the 10 most critical pathways in bladder cancer. We selected the top 660 genes from each method for enrichment analysis (so the number was consistent with the control hub). The larger the bubble in the figure, the greater the overlap between the result of the method and the genes in the pathway.

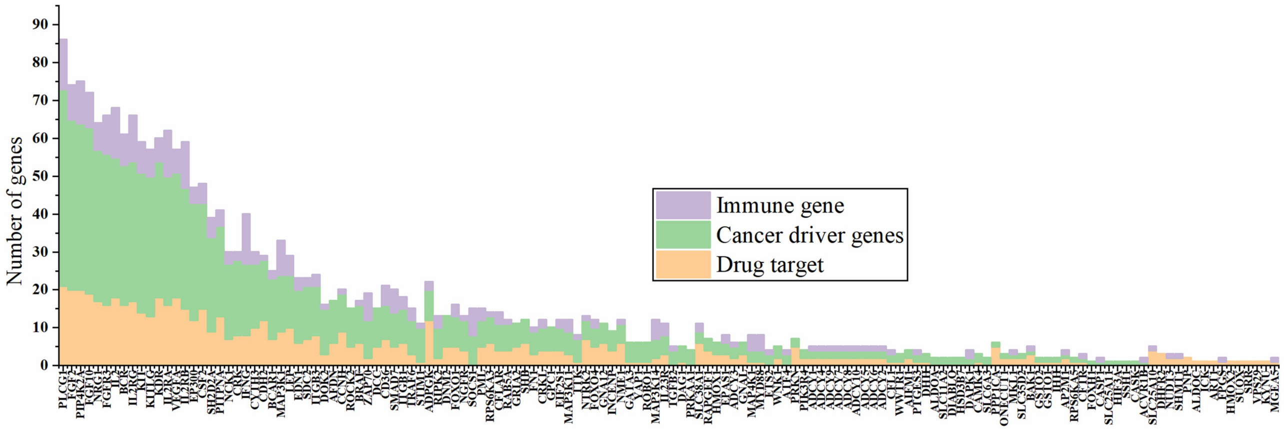


**eFigure 8.** The number of neighbor functional genes of 173 sCKGs. We measured the number of cancer driver genes, drug targets, and immune genes among the neighbors of 173 sCKGs and presented it in the column stacking diagram.

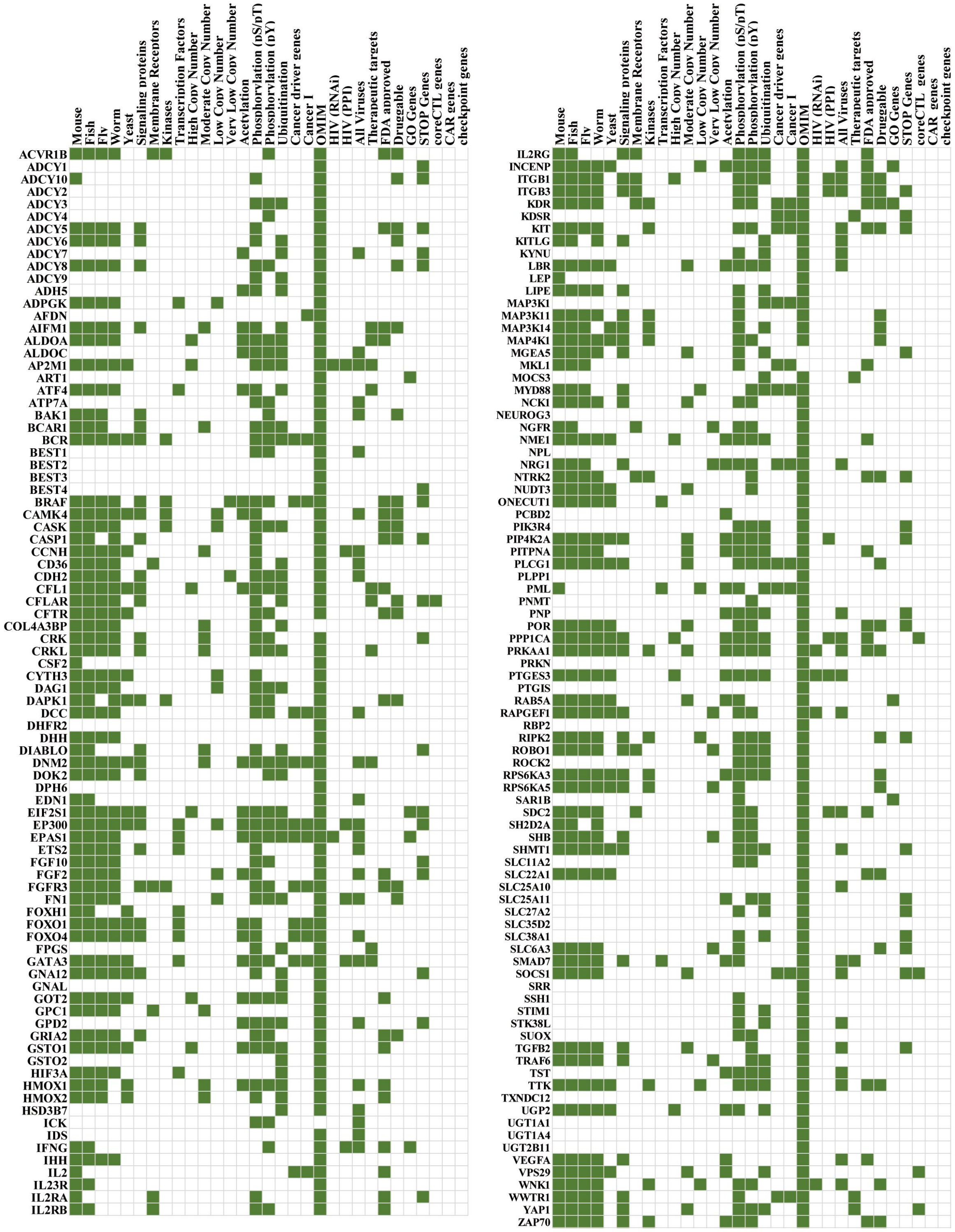


**eFigure 9.** The distribution of 173 sCKGs in different datasets. Each row represents a gene and each column represents a data set. Green squares indicate the overlapping genes in the corresponding dataset.

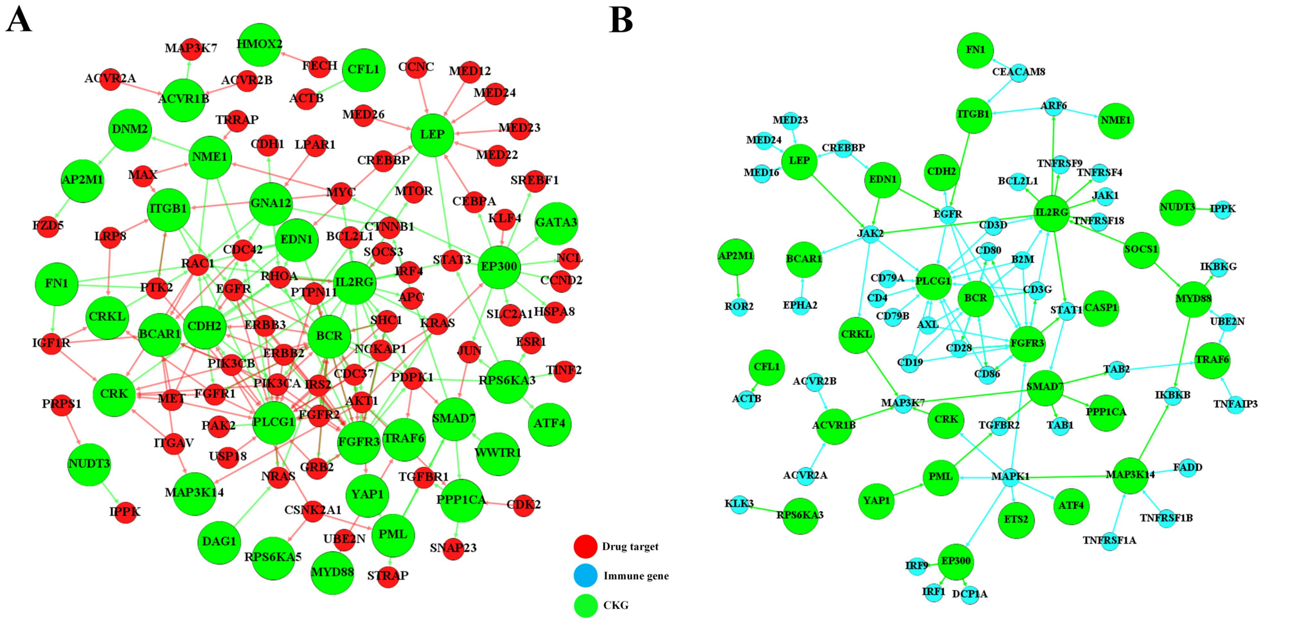


**eFigure 10.** 35 sCKGs and their neighbor functional genes. For the 35 sensitive CKG, we counted the drug target[7] and immune gene[8, 9] contained in its direct neighbors, and the figure shows the network of drug target and immune gene in sCKG neighbors, respectively.

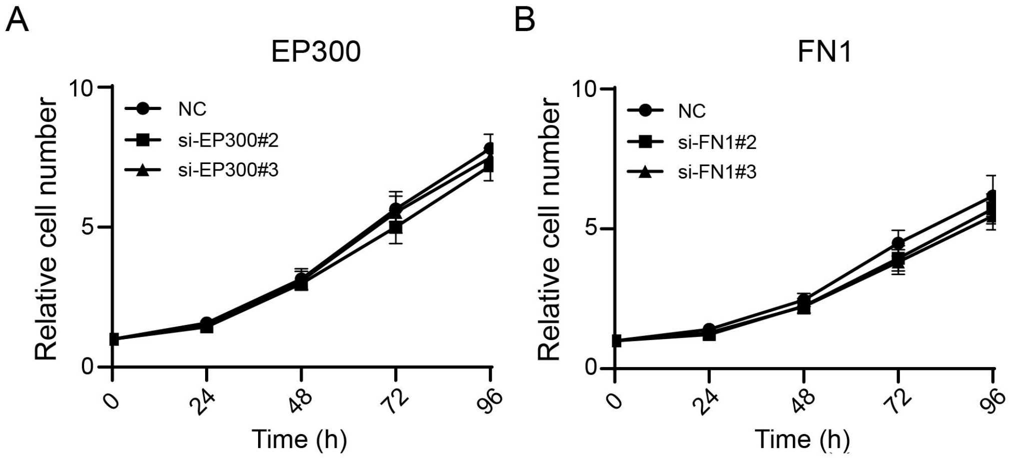


**eFigure 11.** The effects of EP300 (A) and FN1 (B) knockdown by siRNA on the proliferation of T24 bladder cancer cells. CCK 8 assay was used to detect the effects of siRNA knockdown.

**eTable 1. Ten types of control-related interactions**

| Type of interactions | Descriptions |
|---|---|
| *Controls-state-change-of* | first protein controls a reaction that changes the state of the second protein |
| *Controls-transport-of* | first protein controls a reaction that changes the cellular location of the second protein |
| *Controls-phosphorylation-of* | first protein controls a reaction that changes the phosphorylation status of the second protein |
| *Controls-expression-of* | first protein controls a conversion or a template reaction that changes the expression of the second protein |
| *Controls-production-of* | the protein controls a reaction of which the small molecule is an output |
| *Controls-transport-of-chemical* | the protein controls a reaction that changes the cellular location of the small molecule |
| *Chemical-affects* | a small molecule having an effect on the protein state |
| *Catalysis-precedes* | first protein controls a reaction whose output molecule is input to another reaction controlled by the second protein |
| *Consumption-controlled-by* | a directed relation from a small molecule to a protein and the protein is controlling the consumption of the small molecule |
| *Used-to-produce* | a reaction consumes a small molecule to produce another small molecule |

The table shows the ten types of control-related interactions used to build the BLCA_GRN, and each of them represents a type of biomolecular interaction.

**eTable 2. Seed nodes of BLCA gene regulatory network**

| | | | | | | | |
|---|---|---|---|---|---|---|---|
| ARID1A | ASXL2 | ATM | CDKN1A | CDKN2A | CREBBP | CTNNB1 | CUL1 |
| DIAPH2 | ELF3 | EP300 | ERBB2 | ERBB3 | ERCC2 | FAT1 | FBXW7 |
| FGFR3 | FOXA1 | FOXQ1 | GNA13 | HRAS | KANSL1 | KDM6A | KLF5 |
| KMT2C | KMT2D | KRAS | NFE2L2 | NRAS | PIK3CA | PSIP1 | PTEN |
| RB1 | RBM10 | RHOA | RHOB | RXRA | SF1 | SF3B1 | SPTAN1 |
| STAG2 | TP53 | TSC1 | TXNIP | ZFP36L1 | TTN | MUC16 | SYNE1 |
| RYR2 | HMCN1 | MACF1 | FLG | FAT4 | CSMD3 | OBSCN | BIRC6 |
| CUBN | NEB | SYNE2 | DNAH11 | LRP1B | XIRP2 | DNAH5 | AHNAK2 |
| RYR1 | MUC17 | FAT3 | ZFHX4 | ADGRV1 | AKAP9 | ANK2 | KMT2A |
| RYR3 | SPTA1 | ABCA13 | PDE4DIP | MDN1 | | | |

The seed nodes are composed of 45 known BLCA cancer driver genes and 50 top mutated genes of BLCA based on data from the Cancer Genome Atlas (TCGA) together. After removing duplicate genes, seed nodes finally contain 77 related genes.

**eTable 3. Datasets for functional enrichment analysis**

| Dataset | Source | Complied | Overlap |
|---|---|---|---|
| Essential genes[10, 11] | DEG database http://tubic.tju.edu.cn/deg/OGEE database http://ogeedb.embl.de | 16559 | 4455 |
| Evolutionary Conservation[12] | Mouse | 6255 | 3604 |
| | Fish | 5936 | 3480 |
| | Fly | 5001 | 3020 |
| | Worm | 4755 | 2887 |
| | Yeast | 2527 | 1651 |
| Cell Signaling | Signaling proteins[13] https://www.cellsignal.com/pathways/index.htm | 1289 | 891 |
| | Membrane receptors[14] | 1004 | 332 |
| | Protein kinases[15, 16] http://kinase.com/kinbase/index.html | 544 | 316 |
| | Transcription factors[17] | 1732 | 552 |
| Protein abundance[18] | High copy number (>100 000 copies) | 533 | 389 |
| | Moderate copy number (5000–100 000 copies) | 1136 | 700 |
| | Low copy number (500–5000 copies) | 793 | 444 |
| | Very low copy number (<500 copies) | 766 | 407 |
| Posttranslational modification datasets[19] | Acetylation | 3009 | 1644 |
| | Phosphorylation (pS/pT) | 8859 | 4207 |
| | Phosphorylation(pY) | 5666 | 2872 |
| | Ubiqutination | 6244 | 3044 |
| Disease genes | Cancer driver genes[20] | 739 | 501 |
| | Cancer I[21] Cancer Gene Census http://www.sanger.ac.uk/genetics/CGP/Census/ | 723 | 522 |
| | OMIM disease records, all genetic diseases excluding the trait records http://omim.org/ | 16052 | 6716 |
| Virus targets | HIV (RNAi) [22-25] | 1835 | 214 |
| | HIV (PPI) [26, 27] VirusMINT http://mint.bio.uniroma2.it/virusmint/Welcome.do | 880 | 596 |
| | All Viruses[26-29] | 4711 | 2019 |
| Drug targets | Therapeutic[7] targets(CRISPR-Cas9) | 628 | 424 |
| | FDA approved drug target[30] | 982 | 808 |
| | Druggable genome[31] | 1336 | 999 |
| Regulators of cell proliferation[32] | GO Genes | 1127 | 497 |
| | STOP Genes | 3588 | 1200 |
| Immune genes | coreCTL_genes[8] | 182 | 101 |
| | CAR_genes[9] | 129 | 46 |
| | checkpoint_genes | 47 | 19 |

This table shows detailed information on all datasets used in the enrichment analysis, including the source, the number of genes possessed by the dataset (Complied), and the number of genes (Overlap) overlapping the BLCA_GRN.

**eTable 4. The enrichment of the CKGs in in ten important cancer signaling pathways**

| Pathway | CKGs | Proportion | Genes involved in the pathway |
|---|---|---|---|
| Cell Cycle | CDKN1A, CDKN2A, RB1, E2F1, CDK2, CCNE1, CCND1 | 100% | CDKN1A/CDKN1B, CDKN2A/CDKN2B/CDKN2C, CCNE1, RB1, CCND1/CCND2/CCND3, E2F1/E2F2/E2F3, CDK2/CDK4/CDK6 |
| p53 | MDM2, ATM, CHEK2, TP53, RPS6KA3, CDKN2A | 100% | MDM2/MDM4, ATM, CHEK2, TP53, RPS6KA3, CDKN2A |
| TGFβ | ACVR1B, SMAD2, SMAD3, SMAD4 | 80% | TGFBR1/TGFBR2, ACVR2A/ACVR1B, SMAD2, SMAD3, SMAD4 |
| RTK/RAS | EGFR, ERBB2, ERBB3, ERBB4, MET, FGFR1, FGFR3, KIT, NTRK1, NTRK2, JAK2, CBL, ABL1, PTPN11, KRAS, HRAS, NRAS, BRAF, RAC1, MAPK1, MAP2K1, MAP2K2 | 58.30% | EGFR, ERBB2, ERBB3, ERBB4, MET, PDGFRA, FGFR1, FGFR2, FGFR3, FGFR4, KIT, IGF1R, RET, ROS1, ALK, FLT3, NTRK1/NTRK2/NTRK3, JAK2, CBL, ERRFI1, ABL1, SOS1, NF1, RASA1, PTPN11, KRAS, HRAS, NRAS, RIT1, ARAF, BRAF, RAF1, RAC1, MAPK1, MAP2K1, MAP2K2 |
| PI3K | PTEN, INPP4B, AKT1, MTOR, RPTOR, TSC | 46.20% | PTEN, INPP4B, PIK3R2, PIK3CA/PIK3CB, PIK3R1/PIK3R3, PPP2R1A, AKT1/AKT2/AKT3, STK11, TSC1/TSC2, RHEB, RICTOR, MTOR, RPTOR |
| Myc | MYC, MAX, MLXIPL | 27.30% | MYC, MYCN, MYCL, MAX, MGA, MXD1/MXD3/MXD4, MXI1, MNT, MLX, MLXIP, MLXIPL |
| Wnt | GSK3B, CTNNB1, TCF7L2 | 18.80% | WIF1, SFRP1/SFRP2/SFRP3/SFRP4/SFRP5, RNF43, LRP5/LRP6, DKK1/DKK2/DKK3/DKK4, ZNRF3, AXIN1/AXIN2, GSK3B, AMER1, CTNNB1, APC, TCF7, TLE1/TLE2/TLE3/TLE4, TCF7L1/TCF7L2 |
| Notch | NOTCH1, EP300 | 9.50% | JAG2, ARRDC1, FBXW7, CUL1, NOV, NOTCH1, NOTCH2, NOTCH3, NOTCH4, CNTN6, MAML3, KAT2B, HES-X, CREBBP, EP300, HEY-X, DNER, PSEN2, NCOR1/NCOR2, SPEN, KDM5A, |
| Hippo | YAP1 | 6.67% | DCHS1/DCHS2, FAT1/FAT2/FAT3/FAT4, TAOK1/TAOK2/TAOK3, NF2, SAV1, STK3/STK4, LATS1/LATS2, MOB1A/MOB1B, CRB1/CRB2, YAP1, PTPN14, TAZ, CSNK1E/CSNK1D, TEAD2 |
| Nrf2 | - | - | KEAP1, CUL3, NFE2L2 |

**eTable 5. The sensitive edges and their corresponding confidence scores**

| sensitive edge | confidence score | sensitive edge | confidence score | sensitive edge | confidence score |
|---|---|---|---|---|---|
| ADH5_ESD | 99.90% | ATP7A_ATOX1 | 99.90% | CSF1_CSF1R | 99.90% |
| KYNU_KMO | 99.90% | STIM1_ORAI1 | 99.90% | NTN1_DCC | 99.80% |
| LEP_LEPR | 99.80% | OGT_MGEA5 | 99.60% | UGT1A4_UGT1A1 | 99.50% |
| UGT1A1_UGT1A4 | 99.50% | NLRP1_CASP1 | 99.40% | NFS1_MOCS3 | 99.40% |
| AAK1_AP2M1 | 99.30% | MOCS3_MOCS2 | 99.20% | IHH_PTCH2 | 99.00% |
| GPD2_GPD1 | 99.00% | TST_SUOX | 98.90% | PREB_SAR1B | 98.80% |
| CASK_NRXN1 | 98.70% | PRKN_SNCAIP | 98.50% | DRAP1_FOXH1 | 98.40% |
| LNX1_NUMB | 98.30% | EIF2S1_ATF4 | 98.30% | GNA12_AKAP13 | 98.30% |
| DHH_SMO | 98.10% | ALDOC_ALDOA | 98.10% | ALDOA_ALDOC | 98.10% |
| CYP51A1_LBR | 98.10% | CRK_KIDINS220 | 97.70% | CDK8_CCNH | 97.60% |
| LBR_MSMO1 | 97.50% | EZR_FAS | 97.40% | ETHE1_TST | 97.20% |
| HNF1B_ONECUT1 | 97.00% | USP7_FOXO4 | 96.80% | CDH5_PTPRJ | 96.70% |
| CYTH3_ARF5 | 96.60% | KLF4_EP300 | 96.60% | AIFM1_ENDOG | 96.50% |
| RIPK2_BCL10 | 96.50% | NPL_NANS | 96.50% | NME1_DNM2 | 96.40% |
| MKL1_CYR61 | 96.40% | UNC5B_DAPK1 | 96.20% | CDH2_GRIA2 | 96.20% |
| MAP3K14_RELB | 95.60% | PCBD2_QDPR | 95.30% | MAP4K1_CARD11 | 95.10% |
| RBSN_PIK3R4 | 95.00% | BAK1_DIABLO | 94.40% | YAP1_PML | 94.20% |
| DCC_PITPNA | 94.10% | HIF3A_EPAS1 | 94.00% | FGF10_PTF1A | 94.00% |
| WWTR1_SMAD7 | 93.90% | SSH1_CFL2 | 93.90% | CFL1_MKL1 | 93.70% |
| BDNF_SHC3 | 93.20% | CAMKK2_PRKAA1 | 93.10% | DHFR2_MTHFD1L | 92.90% |
| POR_CYP27B1 | 92.90% | LRP1_ITGB3 | 92.80% | GOT2_HOGA1 | 92.10% |
| SDC2_SPP1 | 92.10% | FGF2_F11R | 92.00% | NCK1_TRIO | 91.90% |
| COPS5_EPO | 91.90% | ERF_ETS2 | 91.80% | MYD88_NGFR | 91.50% |
| PLCG1_CAMK4 | 91.20% | LMTK2_PPP1CA | 91.20% | NRG1_DOCK7 | 91.10% |
| SH2D2A_AXL | 90.30% | ESR2_GNAL | 90.30% | PIF1_PTGES3 | 90.20% |
| SHB_ROCK2 | 90.00% | LPAR4_RPS6KA5 | 90.00% | SPTSSA_KDSR | 90.00% |
| RAB5A_CXCR1 | 90.00% | TRAF6_ITGB3B | 90.00% | PSAT1_VPS29 | 90.00% |

| | | P | | | |
|---|---|---|---|---|---|
| CYBRD1_SLC11A2 | 89.20% | CMA1_KITLG | 86.60% | FPGS_SHMT1 | 85.00% |
| GSTO2_GSR | 84.60% | GATA3_IL10 | 83.50% | COL4A3BP_COL4A3 | 83.40% |
| TESK1_CFL1 | 82.90% | SLC38A2_SLC38A1 | 82.20% | CP_HEPH | 81.70% |
| ZAP70_DUSP3 | 81.40% | IFNG_IL23R | 80.20% | DOK2_NCK2 | 77.00% |
| IL2_IL27RA | 76.10% | STK24_STK38L | 71.20% | SLC25A11_MDH2 | 70.60% |
| GSTO1_AS3MT | 66.80% | SC5D_CYP7A1 | 61.50% | FOXH1_GSC | 61.20% |
| BCR_AFDN | 56.40% | CBS_SQOR | 55.80% | NAGLU_IDS | 55.00% |
| NTRK2_KCNA3 | 54.00% | NR1H4_HSD3B7 | 53.60% | USF2_HMOX1 | 51.50% |
| FGFR3_RPS6KA3 | 48.90% | HES6_NEUROG3 | 41.00% | SIAE_NPL | 40.70% |
| UGP2_PPP1R3C | 40.30% | CDK20_ICK | 40.10% | HSD3B7_PTGIS | 38.80% |
| E2F7_APAF1 | 35.80% | PIAS4_SOCS1 | 35.20% | SLC7A10_SRR | 34.60% |
| DHCR7_POR | 33.40% | RBP2_BRD2 | 33.30% | FH_SLC25A11 | 33.00% |
| TGFB2_PTHLH | 31.90% | GPC1_ROBO1 | 31.70% | SLC25A10_MDH1 | 31.60% |
| ICK_BAG6 | 31.30% | FOXO1_CDKN2B | 29.20% | GATA2_EDN1 | 28.60% |
| MAP3K11_PIN1 | 28.20% | PIK3R4_PIP4K2A | 26.60% | ACVR1B_TDP2 | 26.30% |
| FOXO4_ZFAND5 | 25.80% | NUDT3_IPPK | 24.40% | ETS2_KIT | 23.70% |
| CYP27A1_SLC27A2 | 22.20% | TXNDC12_SRXN1 | 18.80% | STK38L_NUAK1 | 18.30% |
| LIPE_MOGAT2 | 17.00% | SLC46A1_DHFR2 | 15.30% | ADCY7_INCENP | - |
| ADPGK_GNPNAT1 | - | DPH6_ART1 | - | PNP_LIPT1 | - |
| BRAF_TTK | - | CD36_TIRAP | - | PLPP1_DAG1 | - |
| SLC22A1_PNMT | - | ATF7_TGFB2 | - | PRPF4B_ELK1 | - |
| MAP3K1_MAFA | - | PNMT_RNLS | - | PTGIS_AKR1D1 | - |
| RPS6KA5_WNK1 | - | SLC35D2_GALT | - | SLC6A3_NDUFV2 | - |
| UGT2B11_ABHD10 | - | | | | |

In eTable5, eTable6, and eTable7, we only list the information in which 35 sCKGs have neighbors as cancer drivers, drug targets, and immune genes.

**eTable 6. Neighbor Cancer driver of the 35 sCKGs**

| 35 sCKGs | Cancer driver | Cancer driver neighbors of 35 sCKGs |
|---|---|---|
| ACVR1B | - | ACVR2A |
| AP2M1 | - | DNM2 |
| ATF4 | - | HERPUD1, DDIT3, MAPK1 |
| BCAR1 | - | HSP90AA1, PIK3CB, SYK, MAP2K4, ITGAV, MET, PIK3CA, PRKCB, PTK6, RAC1, RET, CXCR4, LCK, JAK2, PIK3R1, KDR |
| BCR | yes | FGFR1, ABL1, FGFR4, PIK3CA, ERBB4, NRG1, RHOH, LCK, KIT, STAT5B, ITK, KDR, PIK3R1, VAV1, AKT2, FGFR3, ERBB2, PDGFB, AKT1, ABI1, SYK, PTEN, FES, PTPN11, AKT3, EGFR, CD28, HSP90AA1, PIK3CB, PDGFRB, ERBB3, SRC, FGFR2, PDGFRA, B2M, BTK, PLCG1 |
| CASP1 | - | TP53 |
| CDH2 | - | FGFR1, PIK3CB, ABL1, PIK3CA, ERBB4, RAC1, MET, RHOA, ERBB3, PTPN11, ERBB2, EGFR, CTNND2, KIF5B, PIK3R1, CTNND1 |
| CFL1 | - | SRC, MKL1 |
| CRK | - | NTRK1, ABL1, FGFR1, PIK3CB, MAP2K4, ITGAV, MET, PDGFRB, RHOA, PIK3CA, MAPK1, PRKCB, ABL2, RAC1, RET, PDGFB, SRC, NTRK3, MAP3K1, PIK3R1 |
| CRKL | - | NTRK1, FGFR1, RAC1, NTRK3, MAP3K1, JAK2 |
| DAG1 | - | PRKCB, NRAS, HRAS, NFATC2 |
| DIABLO | - | BIRC3, CASP9 |
| EDN1 | - | HIF1A, COL3A1, RAC1, RHOA, GNAQ, GATA2, ARNT, GNA11, CREBBP, AKT1, EP300, JAK2, EGFR, CYSLTR2 |
| EP300 | yes | RB1, FOXO4, NOTCH1, TERT, PML, HEY1, CCND2, ATM, AR, CDKN1A, DNAJB1, AKT1, KLF4, CEBPA, SMAD4, SMAD2, BAX, GATA1, FOXO3, GATA3, JUN, HIST1H4I, TP63, STAT3, HIST1H3B, MYC, NCOA2, FOXO1, MAPK1, SMAD3, SGK1 |
| ETS2 | - | KIT, MAPK1, CSF1R |
| FGFR3 | yes | PIK3CA, ERBB4, NRG1, LCK, KIT, FGFR1OP, STAT5B, ITK, KDR, PIK3R1, VAV1, AKT2, PRKCB, RAC1, ERBB2, PDGFB, AKT1, BCR, CNTRL, ABI1, CUX1, SYK, RHOA, PTEN, PTPN11, AKT3, EGFR, TRIM24, CD28, HSP90AA1, PIK3CB, KRAS, PDGFRB, ERBB3, MAPK1, SRC, PDGFRA, B2M, BTK, PLCG1 |
| FN1 | - | PRKAR1A, SDC4, SYK, RAC1, SRC, PRKACA |
| GNA12 | - | CTNNB1, RAC1, RHOA, CDH1, CTNND1 |
| IL2RG | - | PIK3CA, APC, LCK, STAT5B, NRAS, JAK2, CCND3, ITK, PIK3R1, HRAS, PRF1, AKT2, PRKCB, RAC1, SOCS1, MAP2K1, STAT6, CBL, IL2, AKT1, MTOR, HLA-A, ABI1, SYK, IRF4, RHOA, AKT3, JAK1, STAT3, PIK3CB, KRAS, MYC, IL7R, MAP2K2, B2M, BTK, SGK1 |
| ITGB1 | - | PRKAR1A, MYC, RAC1, SRC, MKL1, MAX, PRKACA, EGFR, KDR |

| | | |
|---|---|---|
| LEP | - | CEBPA, HIF1A, STAT3, MED12, NCOA2, PPARG, CCNC, PTPN11, ARNT, NCOA1, STAT5B, CREBBP, EP300, JAK2 |
| MAP3K14 | - | ITGAV, MAPK1, IKBKB, BIRC3, AKT1 |
| MYD88 | yes | NFKB2, IKBKB, SOCS1 |
| NME1 | - | DNM2, MYC, RAC1, MAX, TRRAP |
| PLCG1 | yes | NTRK1, ABL1, FGFR1, FGFR4, PIK3CA, ERBB4, CREB1, NRG1, TRA, LCK, KIT, FGFR1OP, NRAS, JAK2, PRKACA, ITK, KDR, PIK3R1, VAV1, MET, AKT2, PRKCB, RAC1, ERBB2, FGFR3, PDGFB, NTRK3, AKT1, BCR, CNTRL, CUX1, CD79A, SYK, RET, PTEN, PTPN11, AKT3, TRB, CD79B, EGFR, TRIM24, CD28, HSP90AA1, PIK3CB, KRAS, PDGFRB, CBLB, ERBB3, SRC, FGFR2, B2M, BTK |
| PML | yes | SMAD2, TGFBR2, MAPK1, EP300, SMAD3, TP53, CHEK2 |
| PPP1CA | - | AKT1 |
| RPS6KA3 | - | STAT3, HIST1H3B, ESR1, JUN, FGFR3, SRC, CIC |
| RPS6KA5 | - | HIST1H3B |
| SMAD7 | - | SMAD4, CTNNB1, SMAD2, TGFBR2, SMAD3, JUN, TFE3, PML, WWTR1 |
| TRAF6 | - | MALT1, TNFAIP3, MAP2K4, ERC1, BCL10, MYD88, AKT1, TP53, CARD11 |
| WWTR1 | yes | LATS1, LATS2 |
| YAP1 | - | ABL1, ERBB4, PML, AKT1, TP63 |

**eTable 7. Neighbor Drug targets of the 35 sCKGs**

| 35 sCKGs | Drug target | Drug target neighbors of 35 sCKGs |
|---|---|---|
| AP2M1 | yes | FZD5,DNM2 |
| ATF4 | yes | - |
| BCAR1 | - | MET,PIK3CB,CDC42,ITGAV,IGF1R,RAC1,PIK3CA |
| BCR | - | ERBB3,AKT1,PIK3CB,GRB2,FGFR1,EGFR,CDC42,ERBB2,IRS2,SHC1,PTPN11,FGFR2,PDPK1,NCKAP1,CDC37,PIK3CA |
| CDH2 | - | CSNK2A1,ERBB3,MET,PIK3CB,FGFR1,EGFR,CDC42,PTPN11,ERBB2,RHOA,RAC1,PIK3CA |
| CFL1 | yes | ACTB |
| CRK | - | MET,PIK3CB,FGFR1,ITGAV,RHOA,IGF1R,RAC1,PIK3CA |
| CRKL | yes | FGFR1,RAC1,IGF1R,LRP8 |
| DAG1 | - | NRAS |
| EDN1 | - | AKT1,EGFR,CDC42,CREBBP,RHOA,RAC1 |
| EP300 | - | AKT1,CCND2,GATA3,NCL,STAT3,JUN,SLC2A1,CEBPA,KLF4,MYC,HSPA8,SREBF1 |
| FGFR3 | - | ERBB3,AKT1,PIK3CB,GRB2,EGFR,ERBB2,PTPN11,IRS2,RHOA,SHC1,RAC1,KRAS,PDPK1,NCKAP1,CDC37,PIK3CA |
| FN1 | - | IGF1R,PTK2,RAC1 |
| GNA12 | - | LPAR1,CDC42,RHOA,CDH1,RAC1,CTNNB1 |
| HMOX2 | - | FECH |
| IL2RG | - | SOCS3,AKT1,STAT3,PIK3CB,MTOR,APC,RHOA,MYC,SHC1,IRF4,KRAS,NRAS,PDPK1,RAC1,NCKAP1,BCL2L1,PIK3CA |
| ITGB1 | - | PTK2,MAX,LRP8,EGFR,MYC,RAC1 |
| LEP | - | STAT3,MED26,MED12,CEBPA,PTPN11,CREBBP,MED22,CCNC,MED24,MED23 |
| MAP3K14 | - | AKT1,ITGAV |
| MYD88 | - | UBE2N |
| NME1 | - | DNM2,MAX,CDC42,MYC,RAC1,TRRAP |
| NUDT3 | - | IPPK,PRPS1 |
| PLCG1 | - | MET,PTK2,GRB2,EGFR,SHC1,NRAS,KRAS,AKT1,IRS2,PTPN11,USP18,PAK2,FGFR2,CDC37,ERBB3,CDC42,RAC1,PIK3CB,FGFR1,ERBB2,PIK3CA |
| PML | - | SMAD7,CSNK2A1,YAP1,STRAP,TGFBR1 |
| PPP1CA | - | SMAD7,AKT1,SNAP23,TGFBR1,CDK2 |
| RPS6KA3 | - | ESR1,STAT3,JUN,ATF4,PDPK1,TINF2 |
| RPS6KA5 | - | CSNK2A1 |
| SMAD7 | yes | JUN,PDPK1,WWTR1,CTNNB1,TGFBR1 |
| TRAF6 | - | UBE2N,PDPK1,AKT1 |
| WWTR1 | yes | SMAD7 |
| YAP1 | yes | AKT1 |

**eTable 8. Neighbor Immune genes of the 35 sCKGs**

| 35 sCKGs | Immune gene | Immune genes neighbors of 35 sCKGs |
|---|---|---|
| ACVR1B | - | MAP3K7 |
| AP2M1 | - | ROR2 |
| ATF4 | - | MAPK1 |
| BCAR1 | - | JAK2, EPHA2 |
| BCR | - | CD19, AXL, CD80, CD28, B2M, EGFR, CD3G, CD86 |
| CASP1 | - | STAT1 |
| CDH2 | - | EGFR |
| CFL1 | - | ACTB |
| CRK | - | MAP3K7, MAPK1 |
| CRKL | - | MAP3K7, JAK2 |
| EDN1 | - | JAK2, EGFR, CREBBP |
| EP300 | - | IRF9, MAPK1, IRF1, DCP1A |
| ETS2 | - | MAPK1 |
| FGFR3 | - | CD19, STAT1, AXL, MAPK1, CD80, CD28, B2M, EGFR, CD3G, CD86 |
| FN1 | - | CEACAM8 |
| IL2RG | - | BCL2L1, STAT1, SOCS1, B2M, ARF6, CD3D, TNFRSF4, TNFRSF18, CD3G, JAK2, TNFRSF9, JAK1 |
| ITGB1 | - | CEACAM8, ARF6, EGFR |
| LEP | - | MED23, CREBBP, MED24, JAK2, MED16 |
| MAP3K14 | - | MAPK1, FADD, TNFRSF1B, TNFRSF1A, IKBKB |
| MYD88 | - | UBE2N, IKBKB, IKBKG, SOCS1 |
| NME1 | - | ARF6 |
| NUDT3 | - | IPPK |
| PLCG1 | - | CD19, CD79A, AXL, CD80, CD79B, CD28, B2M, CD3D, CD4, EGFR, CD3G, JAK2, CD86 |
| PML | - | YAP1, MAPK1, TGFBR2 |
| PPP1CA | yes | - |
| RPS6KA3 | - | KLK3 |
| SMAD7 | - | MAP3K7, STAT1, TAB1, PPP1CA, TAB2, TGFBR2 |
| TRAF6 | - | TNFAIP3, TAB2, UBE2N |
| YAP1 | yes | - |

**eTable 9. Six drug targets and their neighbor genes of BLCA_GRN**

| Drug Targets | Degree | Node Type | Neighbors | Edge type | Degree | Node Type |
|---|---|---|---|---|---|---|
| CTLA4 | 7 | Tail | FOXP3 | controls-expression-of | 5 | Head |
| | | | FYN | controls-state-change-of | 541 | control hub |

| | | | | | | |
|---|---|---|---|---|---|---|
| | | | LCK | controls-state-change-of | 394 | control hub |
| | | | LYN | controls-state-change-of | 139 | control hub |
| | | | NFATC2 | controls-expression-of | 162 | control hub |
| | | | SRC | controls-state-change-of | 514 | control hub |
| | | | YES1 | controls-state-change-of | 106 | control hub |
| TFDP1 | 48 | Head | APAF1 | controls-expression-of | 11 | Tail |
| | | | ATM | controls-state-change-of | 91 | control hub |
| | | | BBC3 | controls-expression-of | 11 | Tail |
| | | | BRCA1 | controls-expression-of | 74 | control hub |
| | | | CASP7 | controls-expression-of | 21 | control hub |
| | | | CAV1 | controls-expression-of | 32 | Head |
| | | | CCNA1 | controls-expression-of | 76 | control hub |
| | | | CCNA2 | controls-expression-of | 90 | control hub |
| | | | CCND3 | controls-expression-of | 44 | control hub |
| | | | CCNE1 | controls-expression-of | 71 | control hub |
| | | | CCNE2 | controls-expression-of | 68 | Tail |
| | | | CDK2 | controls-state-change-of | 334 | control hub |
| | | | CDKN1B | controls-expression-of | 94 | control hub |
| | | | CDKN2A | controls-expression-of | 42 | control hub |
| | | | CDKN2C | controls-expression-of | 11 | Head/Tail |
| | | | CDT1 | controls-expression-of | 9 | Head/Tail |
| | | | CES1 | controls-expression-of | 55 | control hub |
| | | | CES1P1 | controls-expression-of | 3 | Head/Tail |
| | | | CES2 | controls-expression-of | 55 | control hub |
| | | | CES3 | controls-expression-of | 44 | Tail |
| | | | CES4A | controls-expression-of | 55 | control hub |
| | | | CES5A | controls-expression-of | 55 | control hub |
| | | | CHEK1 | controls-state-change-of | 149 | control hub |
| | | | E2F2 | controls-expression-of | 25 | Head |
| | | | HIC1 | controls-expression-of | 3 | Head/Tail |
| | | | MCL1 | controls-expression-of | 11 | Head/Tail |
| | | | MEF2C | controls-state-change-of | 18 | control hub |
| | | | PLAU | controls-expression-of | 32 | Tail |
| | | | PMAIP1 | controls-expression-of | 7 | Tail |
| | | | RB1 | controls-expression-of | 134 | control hub |
| | | | RBBP8 | controls-expression-of | 7 | Head/Tail |
| | | | RRM1 | controls-expression-of | 8 | Head/Tail |
| | | | RRM2B | controls-expression-of | 11 | Head/Tail |
| | | | SERPINE1 | controls-expression-of | 27 | Tail |
| | | | SIRT1 | controls-expression-of | 77 | Head |
| | | | SPI1 | controls-state-change-of | 17 | Head |
| | | | SULT2A1 | controls-expression-of | 20 | Head/Tail |
| | | | SUZ12 | controls-expression-of | 62 | Head/Tail |
| | | | TBP | controls-state-change-of | 27 | control hub |

| | | | | | | |
|---|---|---|---|---|---|---|
| | | | TK2 | controls-expression-of | 2 | Head/Tail |
| | | | TP73 | controls-expression-of | 83 | control hub |
| | | | TRIM28 | controls-state-change-of | 20 | control hub |
| | | | WASF1 | controls-expression-of | 158 | Tail |
| | | | XRCC1 | controls-expression-of | 9 | Head/Tail |
| CD274 | 22 | Head | CD247 | controls-state-change-of | 271 | Tail |
| | | | CD3D | controls-state-change-of | 88 | Tail |
| | | | CD3E | controls-state-change-of | 95 | Tail |
| | | | CD3G | controls-state-change-of | 266 | Tail |
| | | | CD4 | controls-state-change-of | 76 | Tail |
| | | | HLA-DPA1 | controls-state-change-of | 73 | Tail |
| | | | HLA-DPB1 | controls-state-change-of | 73 | Tail |
| | | | HLA-DQA1 | controls-state-change-of | 73 | Tail |
| | | | HLA-DQA2 | controls-state-change-of | 73 | Tail |
| | | | HLA-DQB1 | controls-state-change-of | 73 | Tail |
| | | | HLA-DQB2 | controls-state-change-of | 73 | Tail |
| | | | HLA-DRA | controls-state-change-of | 121 | Tail |
| | | | HLA-DRB1 | controls-state-change-of | 121 | Tail |
| | | | HLA-DRB3 | controls-state-change-of | 73 | Tail |
| | | | HLA-DRB4 | controls-state-change-of | 73 | Tail |
| | | | HLA-DRB5 | controls-state-change-of | 73 | Tail |
| | | | LCK | controls-state-change-of | 394 | control hub |
| | | | TRA | controls-state-change-of | 19 | Tail |
| | | | TRAC | controls-state-change-of | 19 | Tail |
| | | | TRB | controls-state-change-of | 19 | Tail |
| | | | TRBC1 | controls-state-change-of | 19 | Tail |
| | | | TRBV12-3 | controls-state-change-of | 19 | Tail |
| FKBP1A | 92 | control hub | BAMBI | controls-state-change-of | 38 | Head |
| | | | BMP1 | controls-state-change-of | 30 | Head |
| | | | BMP10 | controls-state-change-of | 8 | Head/Tail |
| | | | BMP15 | controls-state-change-of | 8 | Head/Tail |
| | | | BMP2 | controls-state-change-of | 12 | Head/Tail |
| | | | BMP3 | controls-state-change-of | 8 | Head/Tail |
| | | | BMP4 | controls-state-change-of | 10 | Head/Tail |
| | | | BMP5 | controls-state-change-of | 8 | Head/Tail |
| | | | BMP6 | controls-state-change-of | 8 | Head/Tail |
| | | | BMP7 | controls-state-change-of | 7 | Head/Tail |
| | | | BMP8A | controls-state-change-of | 8 | Head/Tail |
| | | | BMP8B | controls-state-change-of | 8 | Head/Tail |
| | | | CAV1 | controls-state-change-of | 32 | Head |
| | | | CCL11 | controls-state-change-of | 24 | Head/Tail |
| | | | CD247 | controls-state-change-of | 271 | Tail |
| | | | CD3D | controls-state-change-of | 88 | Tail |
| | | | CD3E | controls-state-change-of | 95 | Tail |

| | | | | | | |
|---|---|---|---|---|---|---|
| | | | CD3G | controls-state-change-of | 266 | Tail |
| | | | CD4 | controls-state-change-of | 76 | Tail |
| | | | CNGA1 | controls-state-change-of | 245 | Tail |
| | | | CNGA3 | controls-state-change-of | 28 | Tail |
| | | | CNGB1 | controls-state-change-of | 245 | Tail |
| | | | CNGB3 | controls-state-change-of | 28 | Tail |
| | | | CXCL10 | controls-state-change-of | 27 | Head/Tail |
| | | | CXCL11 | controls-state-change-of | 24 | Head/Tail |
| | | | CXCL13 | controls-state-change-of | 22 | Head/Tail |
| | | | CXCL9 | controls-state-change-of | 30 | Head/Tail |
| | | | CXCR3 | controls-state-change-of | 34 | Head/Tail |
| | | | ERBB2 | controls-state-change-of | 272 | control hub |
| | | | ERBB3 | controls-state-change-of | 265 | control hub |
| | | | FLNA | controls-state-change-of | 35 | Head/Tail |
| | | | FZD2 | controls-state-change-of | 72 | Tail |
| | | | GDF1 | controls-state-change-of | 11 | Head/Tail |
| | | | GDF10 | controls-state-change-of | 8 | Head/Tail |
| | | | GDF11 | controls-state-change-of | 8 | Head/Tail |
| | | | GDF15 | controls-state-change-of | 11 | Head/Tail |
| | | | GDF2 | controls-state-change-of | 7 | Head/Tail |
| | | | GDF3 | controls-state-change-of | 8 | Head/Tail |
| | | | GDF5 | controls-state-change-of | 8 | Head/Tail |
| | | | GDF6 | controls-state-change-of | 8 | Head/Tail |
| | | | GDF7 | controls-state-change-of | 8 | Head/Tail |
| | | | GDF9 | controls-state-change-of | 8 | Head/Tail |
| | | | GDNF | controls-state-change-of | 27 | Head/Tail |
| | | | GNAI1 | controls-state-change-of | 388 | control hub |
| | | | GNAI2 | controls-state-change-of | 398 | control hub |
| | | | GNAI3 | controls-state-change-of | 150 | control hub |
| | | | GNAO1 | controls-state-change-of | 106 | Tail |
| | | | GNAZ | controls-state-change-of | 148 | control hub |
| | | | HLA-DRA | controls-state-change-of | 121 | Tail |
| | | | HLA-DRB1 | controls-state-change-of | 121 | Tail |
| | | | IFNG | controls-state-change-of | 114 | sensitive control hub |
| | | | IFNGR1 | controls-state-change-of | 63 | Tail |
| | | | INHBA | controls-state-change-of | 10 | Head/Tail |
| | | | INHBB | controls-state-change-of | 10 | Head/Tail |
| | | | INHBC | controls-state-change-of | 10 | Head/Tail |
| | | | INHBE | controls-state-change-of | 10 | Head/Tail |
| | | | JAK1 | controls-state-change-of | 188 | Tail |
| | | | JAK2 | controls-state-change-of | 221 | control hub |
| | | | LCK | controls-state-change-of | 394 | control hub |
| | | | LEFTY1 | controls-state-change-of | 16 | Head |

| | | | | | | |
|---|---|---|---|---|---|---|
| | | | LEFTY2 | controls-state-change-of | 16 | Head |
| | | | MSTN | controls-state-change-of | 8 | Head/Tail |
| | | | NFATC1 | controls-transport-of | 110 | control hub |
| | | | NFATC2 | controls-transport-of | 162 | control hub |
| | | | NFATC3 | controls-transport-of | 140 | control hub |
| | | | NODAL | controls-state-change-of | 13 | Tail |
| | | | NRG1 | controls-state-change-of | 155 | sensitive control hub |
| | | | ORAI1 | controls-state-change-of | 135 | Tail |
| | | | PF4 | controls-state-change-of | 34 | Head/Tail |
| | | | PML | controls-state-change-of | 21 | sensitive control hub |
| | | | PRKCQ | controls-state-change-of | 192 | control hub |
| | | | SH3BP2 | controls-state-change-of | 25 | Head/Tail |
| | | | SLC24A1 | controls-state-change-of | 423 | control hub |
| | | | SLC24A2 | controls-state-change-of | 331 | control hub |
| | | | SMAD2 | controls-state-change-of | 104 | control hub |
| | | | SMAD3 | controls-state-change-of | 109 | control hub |
| | | | SMAD7 | controls-transport-of | 30 | sensitive control hub |
| | | | SMURF1 | controls-transport-of | 28 | Head |
| | | | SMURF2 | controls-transport-of | 70 | Head |
| | | | TGFB1 | controls-transport-of | 37 | Head/Tail |
| | | | TGFB2 | controls-state-change-of | 14 | sensitive control hub |
| | | | TGFB3 | controls-transport-of | 21 | Head/Tail |
| | | | TGFBR1 | controls-transport-of | 31 | Tail |
| | | | TGFBR2 | controls-transport-of | 27 | Head/Tail |
| | | | TLL1 | controls-state-change-of | 26 | Head |
| | | | TLL2 | controls-state-change-of | 26 | Head |
| | | | TRPC3 | controls-state-change-of | 145 | Head/Tail |
| | | | TRPC6 | controls-state-change-of | 153 | Tail |
| | | | TRPV6 | controls-state-change-of | 152 | Head/Tail |
| | | | WNT5A | controls-state-change-of | 76 | Head/Tail |
| | | | ZAP70 | controls-state-change-of | 64 | sensitive control hub |
| | | | ZFYVE9 | controls-state-change-of | 11 | Head |
| IL12B | 47 | Head | ALOX12B | controls-expression-of | 9 | Head |
| | | | CCL2 | controls-expression-of | 10 | Head/Tail |
| | | | CCL3 | controls-expression-of | 7 | Head/Tail |
| | | | CCL4 | controls-expression-of | 7 | Head/Tail |
| | | | CCR5 | controls-expression-of | 11 | Head/Tail |
| | | | CD3E | controls-expression-of | 95 | Tail |
| | | | CD4 | controls-expression-of | 76 | Tail |
| | | | CXCL1 | controls-expression-of | 6 | Head/Tail |
| | | | CXCL9 | controls-expression-of | 30 | Head/Tail |

| | | | | | | |
|---|---|---|---|---|---|---|
| | | | EOMES | controls-expression-of | 16 | Tail |
| | | | FASLG | controls-expression-of | 41 | Tail |
| | | | FOS | controls-expression-of | 104 | control hub |
| | | | GADD45B | controls-expression-of | 14 | Head/Tail |
| | | | GADD45G | controls-expression-of | 14 | Head/Tail |
| | | | GZMA | controls-expression-of | 6 | Head/Tail |
| | | | GZMB | controls-expression-of | 21 | Tail |
| | | | IFNG | controls-expression-of | 114 | sensitive control hub |
| | | | IL19 | controls-expression-of | 6 | Head/Tail |
| | | | IL1B | controls-expression-of | 23 | Head |
| | | | IL1R1 | controls-expression-of | 12 | Head/Tail |
| | | | IL23R | controls-expression-of | 33 | sensitive control hub |
| | | | IL24 | controls-expression-of | 6 | Head/Tail |
| | | | IL4 | controls-transport-of | 40 | control hub |
| | | | IL6 | controls-expression-of | 40 | Head/Tail |
| | | | ITGA3 | controls-expression-of | 14 | Tail |
| | | | LCK | controls-state-change-of | 394 | control hub |
| | | | MPO | controls-expression-of | 7 | Head/Tail |
| | | | NFKB1 | controls-state-change-of | 136 | control hub |
| | | | NFKB2 | controls-transport-of | 21 | control hub |
| | | | NFKBIA | controls-state-change-of | 51 | control hub |
| | | | NOS2 | controls-expression-of | 101 | control hub |
| | | | PIK3CA | controls-state-change-of | 360 | Tail |
| | | | PIK3CB | controls-state-change-of | 366 | Tail |
| | | | PIK3CD | controls-state-change-of | 366 | Tail |
| | | | PIK3CG | controls-state-change-of | 366 | Tail |
| | | | PIK3R1 | controls-state-change-of | 327 | Tail |
| | | | RELA | controls-state-change-of | 135 | control hub |
| | | | RELB | controls-transport-of | 10 | Tail |
| | | | STAT1 | controls-state-change-of | 129 | control hub |
| | | | STAT3 | controls-state-change-of | 123 | control hub |
| | | | STAT4 | controls-transport-of | 63 | control hub |
| | | | STAT5A | controls-state-change-of | 113 | control hub |
| | | | STAT6 | controls-state-change-of | 71 | control hub |
| | | | TBX21 | controls-expression-of | 9 | Head/Tail |
| | | | TNF | controls-expression-of | 54 | control hub |
| PTGS2 | 22 | Head | ALOX5AP | catalysis-precedes | 12 | Head |
| | | | CYP4F12 | catalysis-precedes | 2 | Head/Tail |
| | | | CYP4F2 | catalysis-precedes | 26 | Tail |
| | | | CYP4F8 | catalysis-precedes | 29 | Tail |
| | | | CYP8B1 | catalysis-precedes | 2 | Head/Tail |
| | | | FAAH | catalysis-precedes | 35 | Head |
| | | | FAAH2 | catalysis-precedes | 35 | Head |

| | | | | | | |
|---|---|---|---|---|---|---|
| | | | FOS | controls-expression-of | 104 | control hub |
| | | | JUN | controls-expression-of | 116 | control hub |
| | | | LTC4S | catalysis-precedes | 11 | Head |
| | | | MAPK1 | controls-expression-of | 397 | control hub |
| | | | MAPK11 | controls-expression-of | 170 | control hub |
| | | | MAPK14 | controls-expression-of | 236 | control hub |
| | | | MAPK3 | controls-expression-of | 388 | control hub |
| | | | MYB | controls-expression-of | 67 | control hub |
| | | | NFATC1 | controls-expression-of | 110 | control hub |
| | | | NFATC2 | controls-expression-of | 162 | control hub |
| | | | NFATC3 | controls-expression-of | 140 | control hub |
| | | | PLA2G4A | catalysis-precedes | 170 | control hub |
| | | | PTGES3 | catalysis-precedes | 12 | sensitive control hub |
| | | | PTGS1 | catalysis-precedes | 22 | Head |

**eTable 10 Sequences of oligonucleotide primers used for qPCR in the study**

| Primer name | Target sequence (5′ to 3′) |
|---|---|
| FN1_F | CAAGCCAGATGTCAGAAGC |
| FN1_R | GGATGGTGCATCAATGGCA |
| RPS6KA3_F | GTGGCAGAAGATGGCTGTG |
| RPS6KA3_R | TGGGTTAATCTCCTCCTCTCC |
| EP300_F | GATGACCCTTCCCAGCCTCAAA |
| EP300_R | GCCAGATGATCTCATGGTGAAGG |
| FGFR3_F | GAGGTGAATGGCAGCAAGGT |
| FGFR3_R | AAGGTGACGTTGTGCAAGGA |
| CASP1_F | TGCCTGTTCCTGTGATGTGG |
| CASP1_R | TGTCCTGGGAAGAGGTAGAAACAT |
| CDH-2_F | CAACTTGCCAGAAAACTCCAGG |
| CDH-2_R | ATGAAACCGGGCTATCTGCTC |
| AP2M1_F | CTCATCTCCCGAGTCTACCGA |
| AP2M1_R | GTTGGACCGCTTAACGTGGA |
| BCAR1_F | CTGCGTGAGGAGACCTACGA |
| BCAR1_R | CAGGAGGAAGCACCCGTTC |

**eTable 11. Sequences of oligonucleotide siRNA or shRNA used in the study**

| siRNA name | Target sequence (5′ to 3′) |
|---|---|
| st-h-CASP1_001 | CACCACTGAAAGAGTGACT |
| st-h-CASP1_002 | TGGAAGACTCATTGAACAT |
| st-h-CASP1_003 | ATATGCCTGTTCCTGTGAT |
| st-h-CDH2_001 | GTAGCTAATCTAACTGTGA |
| st-h-CDH2_002 | GACCATCACTCGGCTTAAT |
| st-h-CDH2_003 | GCTGCAGATCTATTTACTT |
| st-h-EP300_001 | GCACAGAAGTGAATTCTCA |
| st-h-EP300_002 | GTATGAATCTGCAAACAAT |
| st-h-EP300_003 | GGACTGCAGTCTATCATGA |

| | |
|---|---|
| st-h-FN1_001 | GGAAAACACTATCAGATAA |
| st-h-FN1_002 | CTGCGAGAGTAAACCTGAA |
| st-h-FN1_003 | GCCAACCTTTACAGACCTA |
| st-h-RPS6KA3_001 | GGAACGTGATATCTTGGTA |
| st-h-RPS6KA3_002 | GACAGTTGGTGTACATTCA |
| st-h-RPS6KA3_003 | GCAAGAGATGTATACATAA |
| st-h-FGFR3_001 | ACGTGGAGTTCCACTGCAA |
| st-h-FGFR3_002 | TGCACAACCTCGACTACTA |
| st-h-FGFR3_003 | CAAGCACGTGGAGGTGAAT |
| st-h-RPS6KA3_001 | GGAACGTGATATCTTGGTA |
| st-h-RPS6KA3_002 | GACAGTTGGTGTACATTCA |
| st-h-RPS6KA3_003 | GCAAGAGATGTATACATAA |
| st-h-AP2M1_001 | GGAGGCTTATTCATCTATA |
| st-h-AP2M1_002 | GCGAGAGGGTATCAAGTAT |
| st-h-AP2M1_003 | CGACCATGATGTCATCAAA |
| st-h-BCAR1_001 | CCCACAAGCTGGTGTTCAT |
| st-h-BCAR1_002 | GCAGTTTGAACGACTGGAA |
| st-h-BCAR1_003 | GCTGGATGGAGGACTATGA |
| sh-RPS6KA3 | GGAGGAGATTAACCCACAAAC |

**Captions for Tables File1-File2, which are the individual Excel spreadsheet file that includes the raw data or results.**

**Tables File1: The BLCA gene regulatory network (BLCA_GRN).** The network contains 7,030 genes (in Table Node) and 103,360 directed interactions (in Table Edge).

**Tables File2: The detail information of 660 CKGs in BLCA gene regulatory network (BLCA_GRN).** We applied our novel CKG approach to the BLCA gene regulatory network BLCA_GRN and identified 660 CKGs. This file lists the detail information of 660 CKGs in BLCA_GRN, including their alteration frequency and neighbor genes acting as drug target, cancer driver genes and/or immune genes.
